# Direct optogenetic activation of inner hair cells improves the efficiency and fidelity of auditory encoding

**DOI:** 10.64898/2026.09.24.751732

**Authors:** Wei Li, Victor Bordier, Ara Schorscher-Petcu, Jules Lubetzki, Trinity Pirrone, Jérémie Barral

## Abstract

Severe hearing impairment affects approximately 5% of the global population, with a prevalence projected to increase over the coming decades. Cochlear implant devices pose a neuroprosthetic treatment option to allow sound perception and speech comprehension. However, users are limited in the spectrum of auditory information they can regain because electrical stimulation of the auditory nerve generates a broad spread of current and limits the transmission of sound frequency information. Recently, optogenetic stimulation of the cochlear nerve has emerged as a promising alternative for hearing restoration. Because light can be precisely confined in space, optogenetic stimulation of the cochlea permits to activate auditory neurons within smaller tonotopic regions, thereby improving the encoding of sound frequencies compared to electrical stimulation. However, this approach is limited in terms of dynamic range and ability to follow rapidly varying inputs, thus preventing faithful transmission of auditory information. Here, we developed an alternative approach targeting directly sensory inner hair cells with optogenetics. We demonstrate that this approach decreases the amount of light energy necessary to evoke auditory responses. Direct stimulation of hair cells increases the dynamic range of intensities encoded by neurons in subcortical auditory areas and improves the temporal fidelity of optogenetic responses. Finally, a biophysical model of the auditory system suggests that preserving the synapses between inner hair cells and auditory neurons greatly improves the encoding performance of an optical cochlear implant. Together, our results suggest that targeting sensory inner hair cells is a promising approach for hearing restoration through an optical implant.

## INTRODUCTION

Electrical cochlear implants (eCI) have restored partial hearing to more than one million people with profound hearing loss, enabling speech comprehension and substantially improving quality of life (1). These neuroprosthetic devices bypass auditory mechano-transduction by directly stimulating spiral ganglion neurons (SGN), thereby transmitting auditory information to the brain. Despite their success, eCIs provide only limited restoration of hearing because electrical current spreads broadly within the cochlea, resulting in poor spatial selectivity. Although several stimulation strategies have been developed to reduce current spread (2–4), spectral resolution remains fundamentally constrained. In humans, approximately 10-20 electrodes are used to stimulate a cochlea that normally contains around 4,000 inner hair cells (IHC), each providing frequency-specific input to the auditory nerve. Increasing the number of electrodes has yielded little improvement because overlapping electrical fields limit the independence of adjacent stimulation sites (5). Consequently, hearing restoration by eCI remains rudimentary, particularly for spectral discrimination and intensity coding.

Optical cochlear implants (oCI) have emerged as a promising alternative that could overcome these limitations (6). By using light instead of electrical current, oCI offer the potential for much more spatially confined stimulation, thereby improving frequency selectivity and sound perception (7, 8). This strategy relies on optogenetics (9), in which light-sensitive ion channels are genetically expressed in the membrane of target cells. In animal models, optogenetic stimulation of SGN successfully evokes auditory brainstem responses (10, 11), neuronal activity in the inferior colliculus (8, 11) and importantly a behavioral response (12). These findings establish the feasibility of oCI while highlighting the need for efficient, stable, and safe optogenetic expression in SGN (7).

While most studies have targeted SGN, this approach bypasses key components of cochlear processing. In the healthy cochlea, sensory transduction and synaptic transmission at the IHC-SGN synapse shape auditory coding over a wide range of sound intensities and temporal patterns (13). Cochlear amplification lowers auditory thresholds and expands the dynamic range of hearing, enabling the detection of faint yet behaviorally relevant sounds in noisy environments (14). Furthermore, sound intensity is encoded through the graded recruitment of distinct populations of auditory nerve fibers with different spontaneous rates and sensitivities, relying on the ability of IHC to maintain graded activation (15). Because optogenetic stimulation of SGN activates all fiber populations simultaneously, this physiological recruitment is lost, resulting in a compressed dynamic range (8, 16).

Temporal coding also remains a major challenge for SGN-targeted oCI. Under normal conditions, SGN phase-lock to sound waveforms at frequencies up to approximately 2-4 kHz (17–20), with synchronization preserved up to about 1 kHz in higher auditory centers (20–23). In contrast, optogenetically driven SGN lose reliable entrainment at stimulation rates above approximately 200 Hz, even when using fast opsins with improved kinetics (8, 12, 24). These limitations reduce the faithful transmission of acoustic information and may ultimately constrain hearing restoration.

Here, we tested an alternative strategy by targeting optogenetic actuators to sensory inner hair cells rather than SGN. We hypothesized that preserving the native IHC-SGN synapse would maintain physiological recruitment of auditory nerve fibers, thereby improving intensity coding and temporal fidelity. To evaluate this approach, we compared optogenetic stimulation of IHC and SGN by recording neuronal responses in two major subcortical auditory nuclei, the cochlear nucleus (CN) and the inferior colliculus (IC). By quantifying differences in response threshold, dynamic range, and temporal fidelity, we demonstrate that targeting IHC provides substantial advantages for optical hearing restoration.

## RESULTS

### Optogenetic activation of inner hair cells

To test the possibility of optogenetically activating hair cells, we expressed ChR2-H134R in conjunction with the fluorescent reporter tdTomato under the *Atoh1* promoter. Expression was found in both inner (IHC) and outer (OHC) hair cells from the base to the apex of the cochlea (Fig. S1A). Using a generic promoter for hair cells enabled us to perform *in vitro* whole-cell recordings and photoactivation in both cell types (Fig. S1B). Although OHC displayed only slightly higher depolarization (6.8±1.3 mV, mean±SE, n=8; current of about 55 pA) than previously published (25), IHC were depolarized by as much as 26 mV (±0.2 mV, n=5; current of about 455 pA) reaching a peak potential of −38 mV (Fig. S1C). This suggested that light-induced depolarization may be sufficient to generate an auditory response. By comparison, the receptor potential at the auditory threshold has an amplitude of a fraction of mV (26) and the mechano-transduction current recorded from IHC at saturation is about 1 nA (27). These experiments are also in line with published data (25, 28) and suggest that (1) expression of ChR2 is strong in IHC and (2) depolarization of IHC would have an effect *in vivo*.

**Figure S1:**
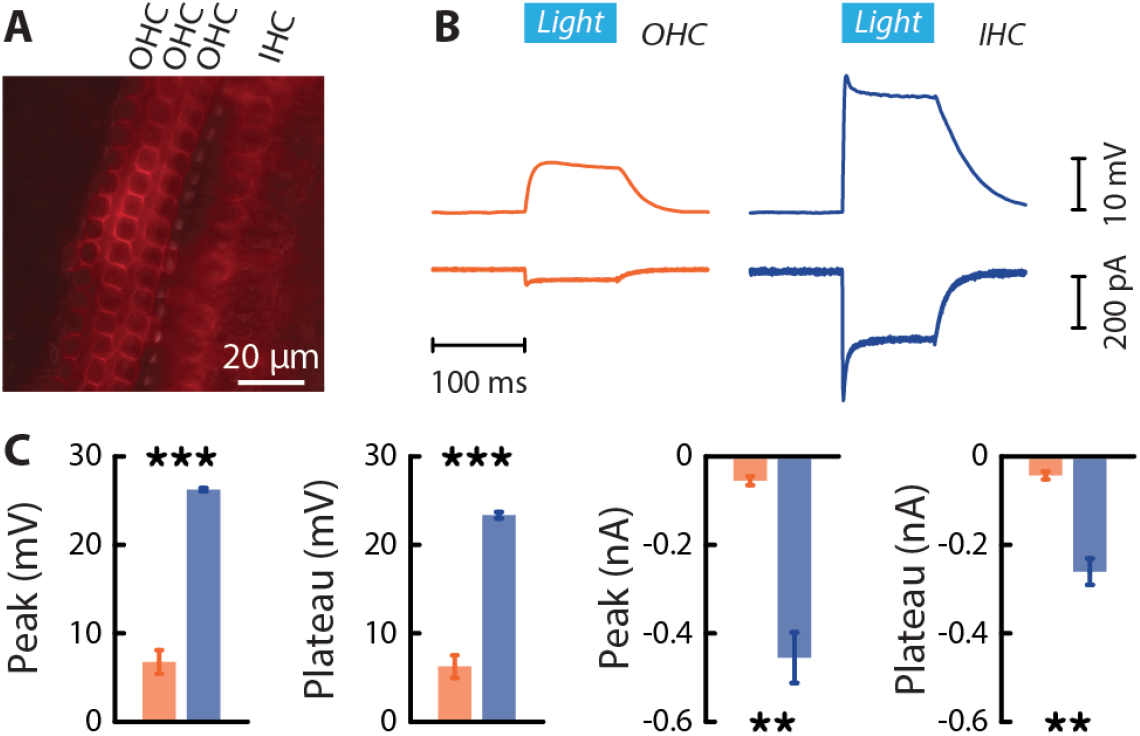
Optogenetic activation of IHC and OHC in vitro. **A.** Superimposed fluorescent and bright field images of the middle turn of the cochlea from a mouse expressing ChR2-tdTomato in IHC and OHC. **B.** Current-clamp (top) and voltage-clamp (bottom) recordings of ChR2 expressing OHC (orange) and IHC (blue) in response to light stimulation (100 ms; 460 nm; 10 mW·mm^−2^). **C.** Peak and plateau potential and current of OHC and IHC upon photostimulation. Cells were obtained from P10-P13 old mice and recorded from the middle turn of the cochlea. Data are presented as mean±SEM (n = 7 OHC, n = 5 IHC). The statistical significance between the two distributions was assessed using a bilateral Student’s T-test (Peak (mV): p = 5.5×10^−6^; Plateau (mV): p = 4.3×10^−6^; Peak (nA): p = 1.9×10^−3^; Plateau (nA): p = 1.2×10^−3^; ** p < 0.01; ***, p < 0.001).

### Optogenetically evoked auditory brainstem response

To assess whether optogenetic stimulation of the cochlea could elicit physiological responses *in vivo*, we used three different mouse lines where opsins were confined to either IHC (Cre expression under the *Myo15a* promoter, ref. (29)), or to SGN (Cre expression under the *Bhlhb5* promoter, ref. (30)), or expressed in both cell types (Cre expression under the *Vglut3* promoter). Fluorescence imaging in explanted cochleae confirmed a robust expression profile restricted to the desired cell types in the cochlea with the appropriate cell membrane localization (Fig. S2).

**Figure S2:**
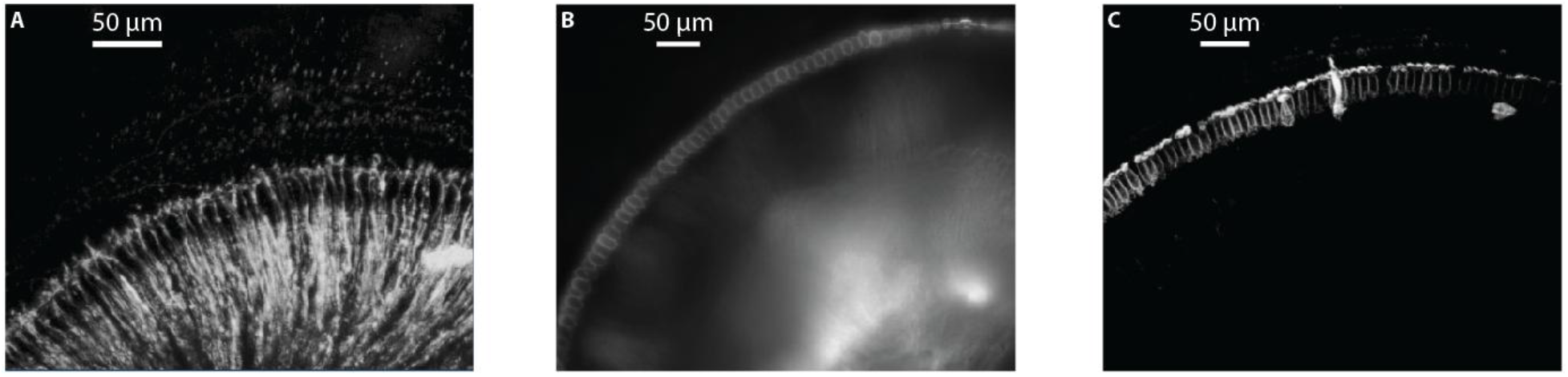
Expression of channelrhodopsin in the cochlea. Expression of ChR2 either in SGN (**A**: Bhlhb5^Cre^::Ai27D, 2-photon fluorescent microscopy), in both SGN and IHC (**B**:Vglut3^Cre^::Ai27D, epifluorescence microscopy), or solely in IHC (**C** Myo15a^Cre^::Ai27D, 2-photon fluorescent microscopy). Note that the expression is precisely localized to the desired cell type in the cochlea but could potentially extend to other cell types in the central nervous system.

Using Auditory Brainstem Response (ABR) measurements, we assessed the efficacy of IHC or SGN optical stimulation to generate evoked potentials. To do this, we surgically ablated the outer and middle ear under isoflurane anesthesia, leaving the stapes in place and approached an optic fiber to the surface of the bony cochlea without cochleostomy (Fig. 1A). Stimulating with a 1 ms light pulse of varying intensity, we obtained robust responses with all three cell targets (Fig. 1B). We found that optogenetic stimulation of SGN generated a first ABR wave, corresponding to synchronized activity in the auditory nerve, earlier than when stimulating IHC or both cell types together (Fig. 1C, top). This was expected given the synaptic delay between IHC and SGN. Regarding the amplitude of the response, no statistical difference was found in terms of absolute value (Fig. 1C, bottom) or growth rate as a function of laser intensity (Fig. S3A). Interestingly, optogenetically evoked ABR did not show signs of decline with age and could be observed in mice as old as 70 weeks (Fig. S3B).

**Figure 1:**
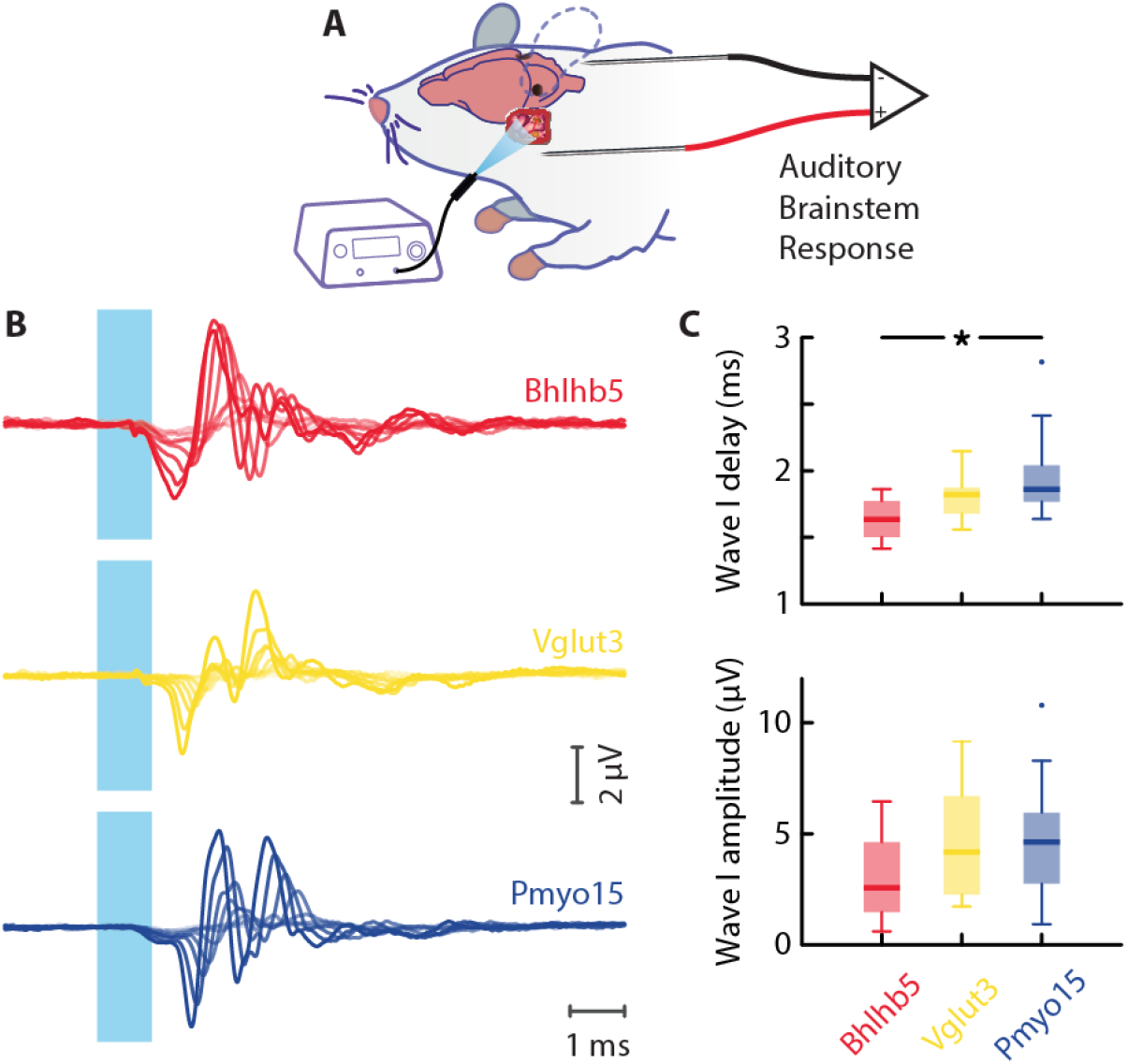
Optogenetically evoked Auditory Brainstem Response. **A.** Experimental strategy to activate the auditory system with artificial optogenetic stimulation using an optic fiber affixed to the cochlea while monitoring the optogenetic Auditory Brainstem Response (oABR). **B.** Examples of oABR in the Bhlhb5^cre^::Ai27D (expression in SGN, red), in the Vglut3^cre^::Ai27D (expression in both IHC and SGN, yellow), or in the Myo15^cre^::Ai27D (expression in IHC, blue) mouse lines. **C.** Wave I delay (top) and amplitude (bottom) for the different mouse lines. Box plots indicate median and interquartile range, whiskers cover the full range of the distribution and outliers are plotted individually (Bhlhb5^cre^::Ai27D, n = 4; Vglut3^cre^::Ai27D, n = 15; Pmyo15^cre^::Ai27D, n = 23). The statistical significance between the two distributions was assessed using a bilateral Mann-Whitney’s U-test (*, p < 0.05; for Wave I delay: Bhlhb5 vs Vglut3: p = 0.26, Bhlhb5 vs Pmyo15: p = 0.04, Vglut3 vs Pmyo15: p = 0.15; for Wave I amplitude: Bhlhb5 vs Vglut3: p = 0.36, Bhlhb5 vs Pmyo15: p = 0.29, Vglut3 vs Pmyo15: p = 0.83).

**Figure S3:**
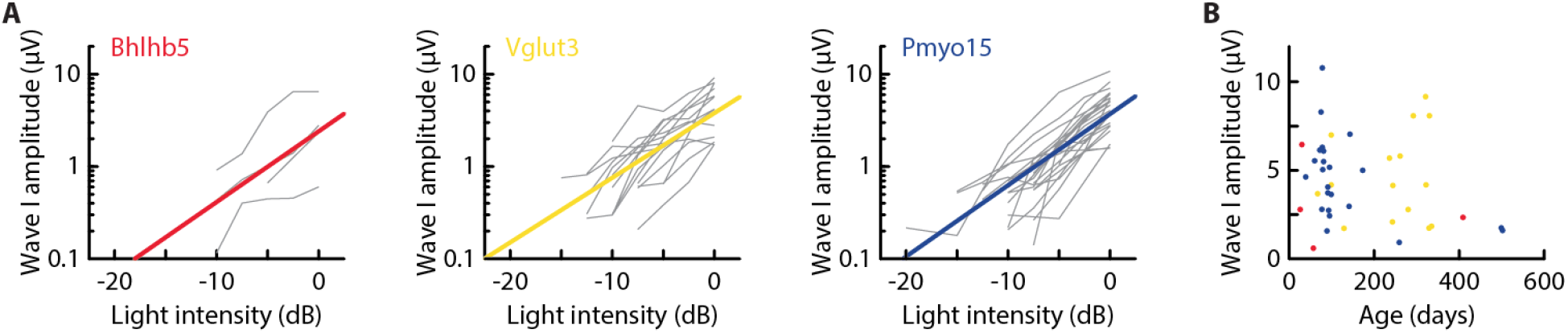
Optogenetically evoked Auditory Brainstem Response as a function of light intensity or age. **A.** Wave I amplitude as a function of light intensity for the different mouse lines (Bhlhb5^cre^::Ai27D, n = 4, red; Vglut3^cre^::Ai27D,n = 15, yellow; Pmyo15^cre^::Ai27D, n = 23, blue). **A.** Wave I amplitude as a function of the age of the animal (same color code as in **A**).

### Optogenetically-evoked subcortical neuronal responses

Having shown that optogenetic stimulation of IHC and/or SGN triggered robust compound brainstem responses, we set out to characterize these at the cellular level in two major nuclei of the auditory pathway. We used high-density extracellular recording probes (Neuropixels) to simultaneously target the ipsilateral cochlear nucleus (CN), which receives direct input from the auditory nerve of the stimulated ear, and the contralateral inferior colliculus (IC), which receives input via both direct and indirect pathways from the CN (Fig. 2A-B). We recorded single- and multi-unit activity from both structures following optogenetic activation of inner hair cells or spiral ganglion neurons (Fig. 2A).

**Figure 2:**
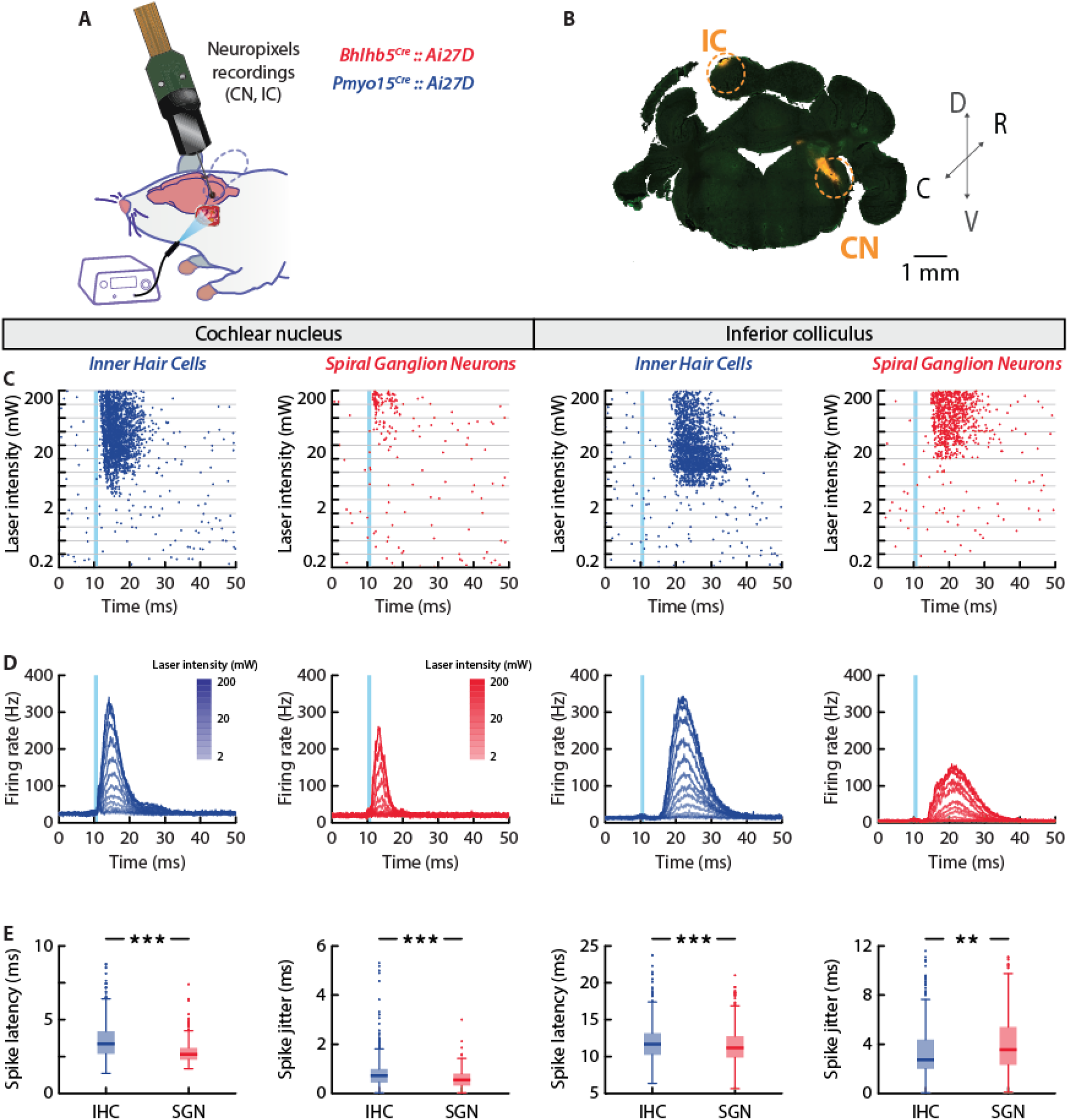
Optogenetically-evoked subcortical neuronal responses. **A.** Optogenetic stimulation of the cochlea combined with simultaneous large-scale neuronal recordings in the cochlear nucleus (CN) and in the inferior colliculus (IC). **B.** Coronal brain slice showing the electrode track targeting the two regions of interest. **C.** Here and thereafter: left half: results from the CN, right half: results from IC. Activation sites are denoted by color, blue: IHC-driven, red: SGN-driven. Exemplar units in the CN (left) and in the IC (right) upon optogenetic stimulation of IHC (blue) or SGN (red). Raster plots displaying 50 trials of 13 light pulse intensities from 0.2 to 200 mW. **D.** Population averages of evoked firing rates (IHC stimulation: n = 630 units in CN, n = 445 units in IC; SGN stimulation: n = 335 units in CN, n = 438 units in IC; only light intensities from 2 to 200 mW are represented). **E**. Latencies of the first optically triggered spike after IHC or SGN stimulation (same n as in **D**). Box plots indicate median and interquartile range, whiskers cover the full range of the distribution and outliers are plotted individually. The statistical significance between the two distributions was assessed using a bilateral Student’s T-test (latency in CN: p = 10^−17^, jitter in CN: p = 6·10^−5^, latency in IC: p = 10^−9^, jitter in IC: p = 2.8·10^−3^; ** p < 0.01; ***, p < 0.001).

Consistent with oABR recordings, the first spikes evoked by optogenetic activation of IHC exhibited a longer latency compared to those triggered by activation of SGN (3.6±0.2 ms for IHC activation vs 2.9±0.2 ms for SGN activation in the CN, mean±SEM), reflecting the additional synapse along the pathway (Fig. 2C and E). Spike timing was also slightly less precise under these conditions (Fig. 2E, 0.84±0.04 ms for IHC activation vs 0.57±0.03 ms for SGN activation in the CN, mean±SEM).

Despite brief stimulation, neural activity was not limited to a single spike in either structure. This was particularly clear in the IC, where we observed sustained activation lasting up to ∼20 ms beyond stimulus onset (Fig. 2D). We quantified the duration of the response to a single light pulse as the decay time constant of the autocorrelation function (Fig. S4). This characteristic timescale increased with increased light intensity and was substantially larger upon IHC stimulation in the CN whereas timescales were more comparable in the IC at high intensities.

**Figure S4:**
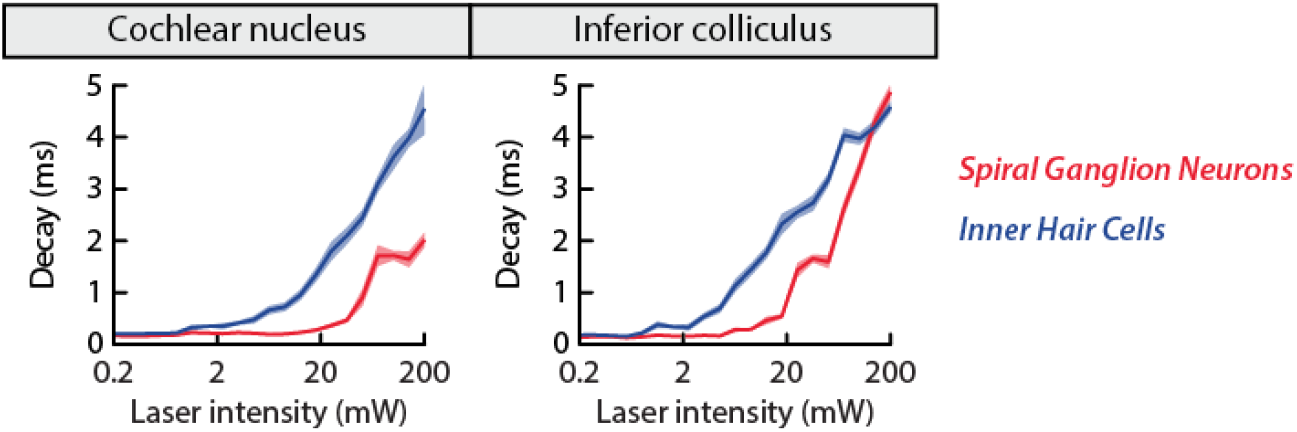
Timescale of responses to optogenetic stimulation. Characteristic timescale, measured as the decay time constant of the autocorrelation function, for neuronal activity in the CN (left) and in the IC (right) upon optogenetic stimulation of IHC (blue) or SGN (red).

### Dynamic range of subcortical neuronal responses to optogenetic stimulation

During this sustained period of activity, neurons showed an intensity-dependent increase in firing rate, typically in a monotonic fashion (Fig. 3A). Activation of sensory hair cells elicited stronger responses in both the CN and IC. Nonmonotonic response functions were observed in 10% of CN neurons and 30% of IC neurons, similar to results obtained with acoustic stimulation in the CN (31, 32) or in the IC (33, 34). In contrast, when SGN were activated, these values decreased to 5% and 10%, respectively, likely due to an inability to achieve maximal activation even with light intensities up to 200 mW.

**Figure 3:**
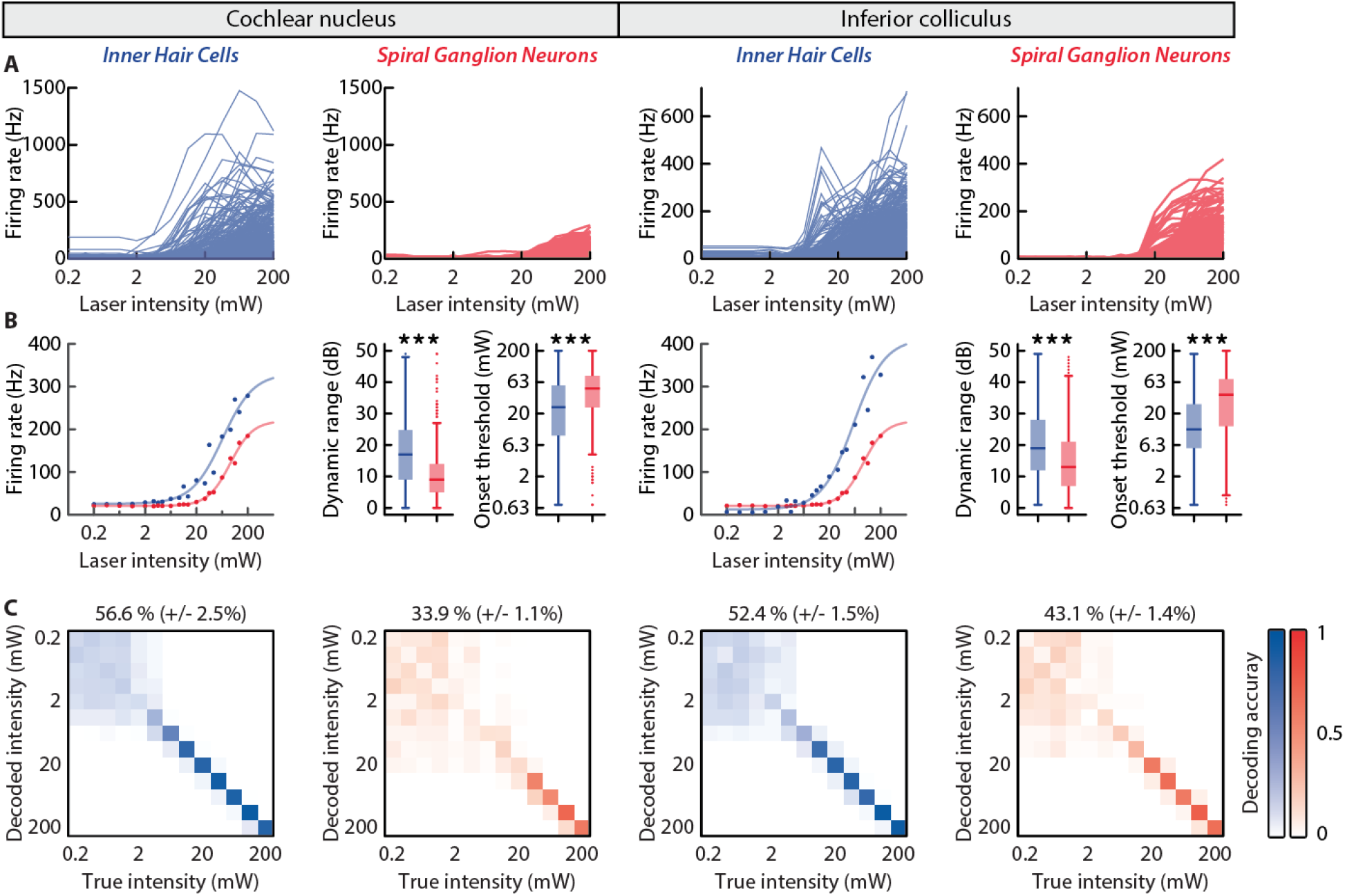
Dynamic range of subcortical neuronal responses to optogenetic stimulation. Left half: results from the CN. Right half: results from the IC. Activation sites are denoted by color, blue: IHC-driven, red: SGN-driven. **A.** Single unit firing rate as a function of laser intensity. **B.** Population average of firing activity as a function of laser intensity fitted with a sigmoid curve (left). Dynamic range (middle) and onset threshold (right) of single units. Box plots indicate median and interquartile range, whiskers cover the full range of the distribution and outliers are plotted individually (IHC stimulation: n = 630 units in CN, n = 445 units in IC; SGN stimulation: n = 335 units in CN, n = 438 units in IC). The statistical significance between the two distributions was assessed using a bilateral Student’s T-test (p < 10^−10^ for any comparison between IHC and SGN stimulation; ***, p < 0.001). **C.** Confusion matrices resulting from decoding light intensity with a SVM classifier and using 256 neurons in each condition out of the total neuronal pool. Average decoding accuracy is mentioned above each matrix (mean±STD, n = 100 random selections of 256 neurons, p < 10^−10^ for any comparison between IHC and SGN stimulation).

The response function width, i.e. the range of laser intensities over which the response amplitude varies, is an important factor for the ability of the neurons to encode intensity information and is denoted as the dynamic range of sensitivity. In the CN, the response function width from 10% to 90% of maximal response was significantly larger for IHC than for SGN stimulation, with average values for single units of 17±0.7 dB and 11.5±0.7 dB, respectively (mean±SEM). Larger values (20.3±1.0 dB and 15.6±0.8 dB) were found in the IC, suggesting that the architecture of the network between these two brain structures and/or intrinsic properties of neurons permit to enlarge the dynamic range as the input propagates through the auditory pathway. The activation threshold was also lower when stimulating IHC. In the IC, neuronal responses were observed with light intensities as low as 11.5±4.2 mW for IHC activation, compared to 27.5±11.3 mW for SGN activation (Fig. 3B). Thresholds were slightly higher in the CN (18.9±7.3 mW and 39.6±14.4 mW for IHC and SGN activation, respectively) but again the threshold was more than doubled from SGN to IHC stimulation target.

Because single neuron measurements are noisy, we also evaluated their population counterparts. To do so, we averaged the firing rates for each light intensity and fitted a sigmoid function (Fig. 3B). The quality of the fit permitted to estimate its parameters with higher confidence than for single neurons. Therefore, we chose to estimate the width (10% - 90%) of the response curve from these fitting parameters. We found a dynamic range of 26±14 dB (27±13 dB) in the CN (IC) upon IHC stimulation and 19±5 dB (14±6 dB) in the CN (IC) upon SGN stimulation. Similarly, the threshold, defined as an increase of the sigmoid by 10%, was increased from 14.2±2.3 mW in CN (12.4±2.3 mW in IC) upon IHC stimulation to 30.3±1.0 mW in CN (30.6±3.8 mW in IC) upon IHC stimulation. Taken together, IHC stimulation provided a 50% increase of dynamic range and permitted to reduce the light intensity by half.

For IHC stimulation, response function centers were distributed across a broader range of intensities, suggesting that population of neurons could encode a wider range of intensities than when SGN were activated. To demonstrate that expanded dynamic range and distribution of sensitivities could enhance coding capacity, we approximated the ability of the neuronal population to encode intensity levels using a Support Vector Machine (SVM) algorithm. The resulting confusion matrix (Fig. 3C) allowed us to estimate the average decoding accuracy. To account for the different number of units per condition, we trained the decoder on various numbers of units and found that the decoding accuracy increased with population size and plateaued after ∼200 neurons (Fig. S5). Therefore, we fixed the population to 256 randomly selected neurons for each region and stimulation mode in subsequent analysis. With this number of neurons, the decoding accuracy was higher for IHC stimulation (56.6% for CN, 52.4% for IC) compared to SGN stimulation (33.9% in CN, 43.1% in IC). These findings demonstrate that valuable information was preserved in spike rate and that pooling responses from neurons sensitive to different intensity ranges improved decoding accuracy. We also computed the channel capacity, which indicates the number of independent levels of a stimulus that can be faithfully transmitted. When converted to decibel units, these measures showed higher values for IHC stimulation (27.4 dB for CN, 25.8 dB for IC) than for SGN stimulation (17.2 dB for CN, 20.6 dB for IC), again demonstrating an increase of 5-10 dB in dynamic range.

In summary, optogenetic stimulation of IHC resulted in lower activation thresholds and a broader dynamic range, as evidenced by the spiking activity recorded from both the CN and the IC and by the information content carried by these structures.

**Figure S5:**
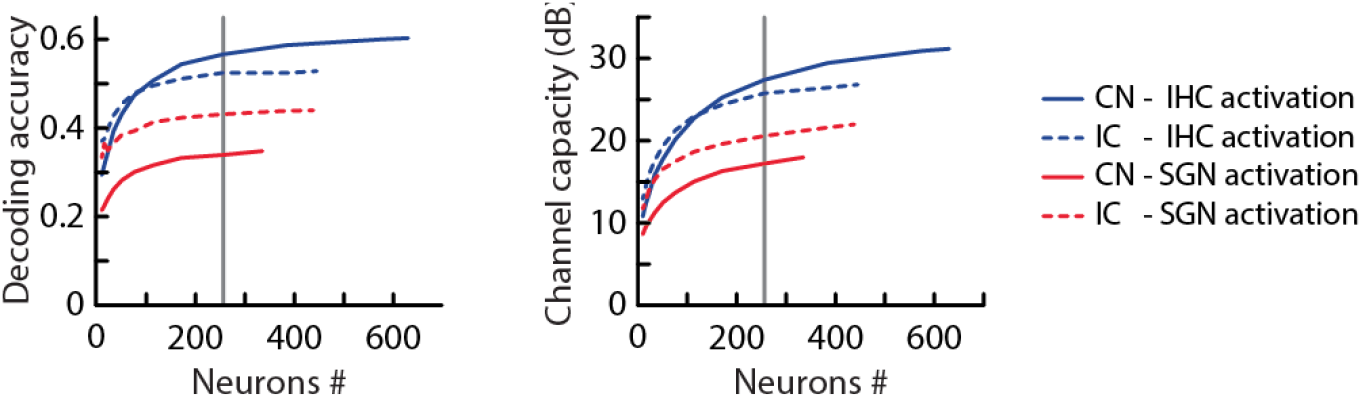
Influence of neuron numbers on the ability to decode the light intensity of a single pulse. Left: Decoding accuracy of light intensity as a function of unit number for CN neurons (solid line; blue for IHC activation, red for SGN activation) and for IC neurons (dashed line; same color code). Right: channel capacity converted to decibels (same line and color code). Data resulted from 100 random selections of various numbers of neurons from the whole population (IHC stimulation: n = 630 units in CN, n = 445 units in IC; SGN stimulation: n = 335 units in CN, n = 438 units in IC). Note that the channel capacity is bound to 60 dB which represents the range of intensities that were used during experiments. The grey line denotes a population of 256 neurons, whose confusion matrices are shown in the main text.

### Entrainment of subcortical neuronal activity by trains of light pulses

To test the ability of the auditory system to follow fluctuating inputs, we used trains of 1 ms light pulses of different intensities (35, 65, and 200 mW) and repetition rates (from 20 to 200 Hz) and again measured neuronal activity in the CN and the IC (Fig. 4A). As shown in this example, IHC stimulation produced more sustained and synchronized responses that we further quantified. At the level of the population, the peristimulus time histogram (PSTH) averaged across all units displayed larger amplitudes and stronger phase-locked modulations (Fig. 4B and Fig. S6).

**Figure 4:**
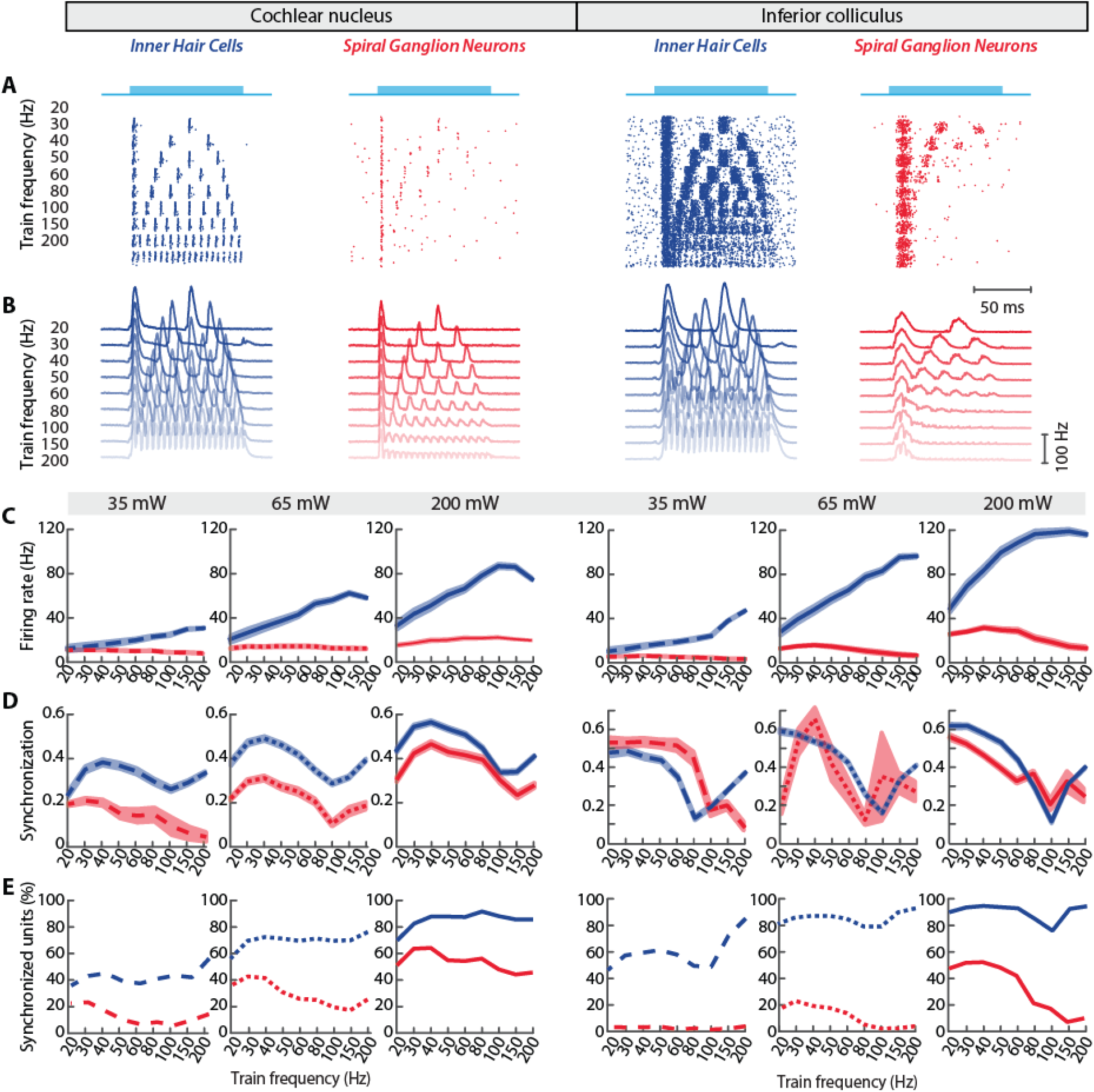
Entrainment of subcortical neuronal activity by trains of light pulses. Left half: results from the CN. Right half: results from the IC. Activation sites are denoted by color, blue: IHC-driven, red: SGN-driven. **A.** Exemplar units in the CN (left) IC (right) upon optogenetic stimulation of IHC (blue, top) or SGN (red, bottom). Raster plots displaying 50 trials of various train frequencies. **B.** Population averages of evoked firing rates for light intensities of 200 mW (IHC stimulation: n = 630 units in CN, n = 445 units in IC; SGN stimulation: n = 335 units in CN, n = 438 units in IC). **C.** Mean (±SEM) evoked firing rate during the stimulus duration as a function of pulse rate for three laser intensities (35 mW, 65 mW, 200 mW). **D.** Mean (±SEM) spike synchronization index of statistically synchronized units (Rayleigh test, p < 0.01) to laser pulses as a function of pulse rate. Laser intensity as in **C**. **E**. Proportion of significantly phase-locked units as a function of pulse rate. Laser intensity as in **C**.

To quantify the sustained response, we measured the evoked firing rate during the whole duration of the stimulus (Fig. 4C). If neurons were able to follow each pulse during the train sequence, increasing the pulse frequency should increase the overall firing rate, simply because of the greater numbers of light pulses. With IHC stimulation, we observed a gradual increase of firing rate up to ∼100 Hz in both brain structures (Fig. 4C). This relationship was observed at every stimulus intensity, but the slope depended on the actual light power. Conversely, increasing the frequency of pulse trains had either no effect or decreased the firing rate in the case of SGN stimulation.

To characterize the ability of neurons to follow fast temporal inputs, we measured the degree of synchronization between spikes and pulse patterns (Fig. 4D). Synchronization was more efficient with IHC stimulation with synchronization indices up to 0.55 (0.65) in the CN (IC). In this case, more than 90% of neurons showed significant synchronization in either brain structure at high light intensities (Fig. 4E). These values were strongly reduced after SGN activation, both in terms of absolute value (Fig. 4D) or proportion of synchronized units (Fig. 4E). We conclude that optogenetic stimulation of IHC results in a higher ability to follow temporally modulated inputs.

**Figure S6:**
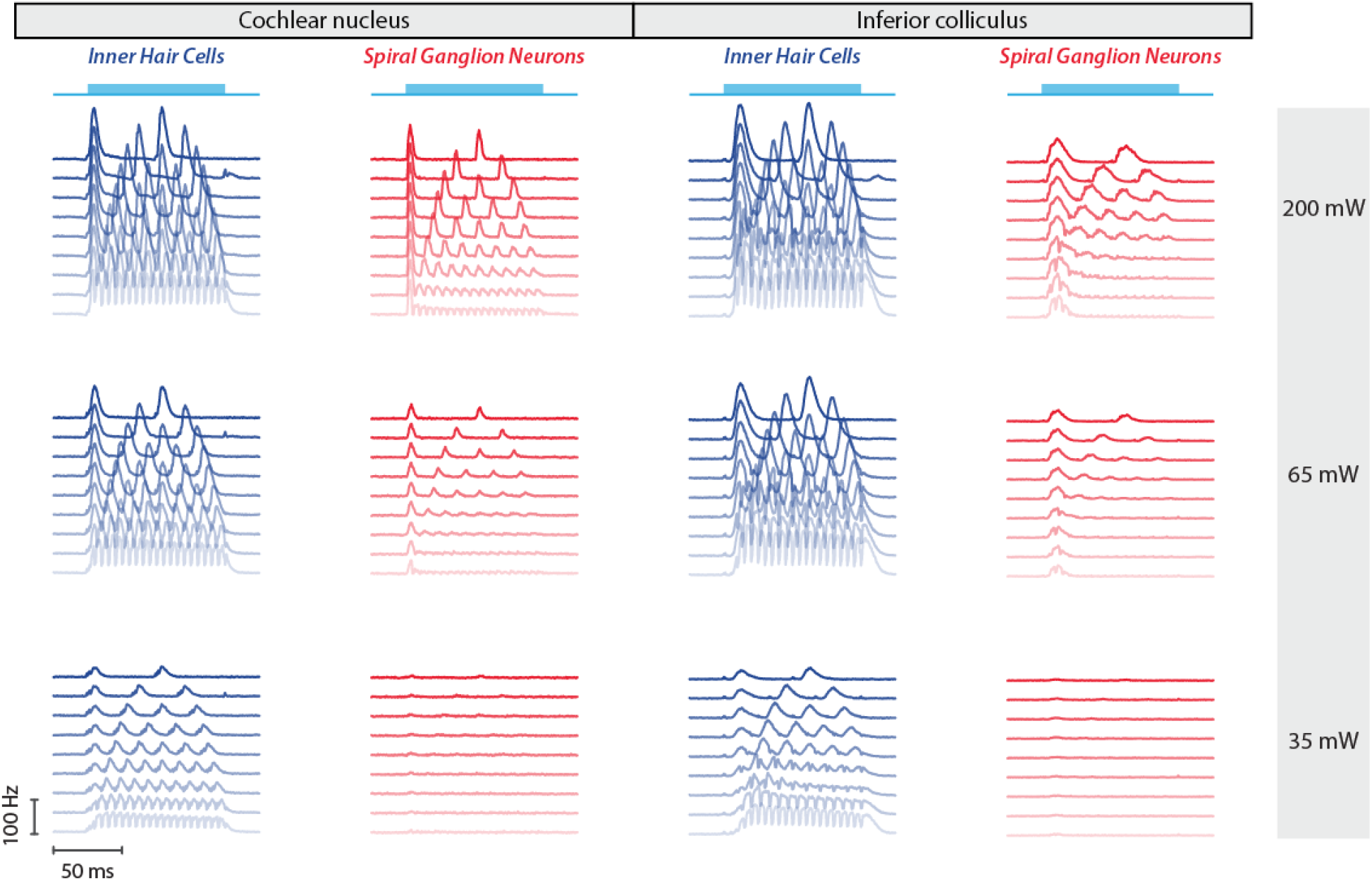
Population firing rate upon pulse train stimulation. Left half: results from the CN. Right half: results from the IC. Activation sites are denoted by color, blue: IHC-driven, red: SGN-driven. Population averages of evoked firing rates for light intensities of 35, 65, and 200 mW at different activation sites. The train frequency varied from 20 Hz (dark shade) to 200 Hz (light shade) (IHC stimulation: n = 630 units in CN, n = 445 units in IC; SGN stimulation: n = 335 units in CN, n = 438 units in IC).

When activated by sounds, neurons in the CN and the IC exhibit characteristic firing patterns. For example, at large sound intensity, octopus and On-L cells maintain a precise phase locking to amplitude fluctuations, whereas other cells produce more sustained responses. However, it is difficult to assess the cell type with extracellular recordings, and some recordings could also result from multiunit activity. To determine whether optogenetic stimulation nonetheless evoked distinct firing patterns in the two stimulation paradigms, we applied Principal Component Analysis (PCA) to the ensemble response profiles, separately for each brain region but by concatenating data from both stimulation modes (Fig. S7). Whereas the first principal component accounted for ∼27% (32 %) of the variance in CN (IC) and showed rather slow modulations, the second component (∼7% of explained variance in both areas) displayed a rapid dynamic, reminiscent of the fast neuronal activity that is prominent in certain CN neurons. The third component (∼5% of the variance) exhibited a dual peak that could be related to neurons firing in bursts. Interestingly, the contribution of units from SGN stimulation was markedly smaller for the fastsecond component, suggesting this stimulation mode was less adapted to follow fast stimulus statistics.

**Figure S7:**
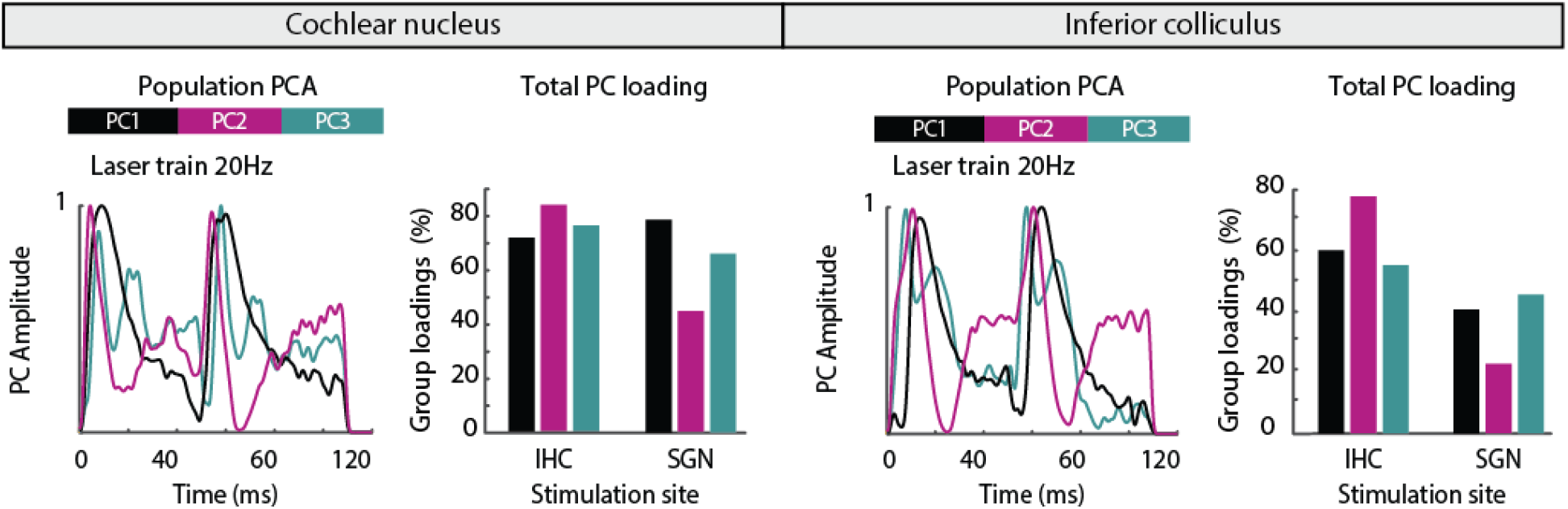
Principal component analysis of the neuronal dynamics upon pulse train stimulation. **A.** Principal component analysis of all recorded units. PCA analysis was performed on both experimental groups (IHC-driven and SGN-driven) simultaneously but separately for each recording region (CN and IC). The first three principal components are shown and PC trajectories plotted as function of time from 0 to 120 ms. Right: Proportion of PC loadings contributed by each experimental group, which indicates the predominance of data originating from IHC stimulation in the PCA analysis, especially for the fast second component.

### Population-level encoding of stimulation frequencies

The synchronization index measures phase locking of individual neurons. To assess the ability of the whole neuronal population to track fast temporal fluctuations, we computed similarity matrices of neuronal response vectors (Fig. 5A). Each value in this matrix represents an estimate of the cross-correlation between PSTH of each neuron. A high correlation value between two response vectors indicates that the same neurons responded in a similar fashion to different stimuli. Because we considered here the correlation between PSTH, we expected to find high correlation values between stimuli of same train frequency but different light intensities. These diagonal elements in off-diagonal blocks were obvious with IHC stimulation but did not stand out clearly with SGN stimulation (Fig. 5A). Instead, we observed a block structure indicating that the neuronal population responded similarly to a given light intensity regardless of pulse-train frequency. This suggests that pulse train frequency may not be well recoverable from neuronal activity. Conversely, IHC activation generated response vectors with strong correlations for shared pulse train frequencies, indicating that the population could phase lock to the light patterns for a large range of frequencies.

**Figure 5:**
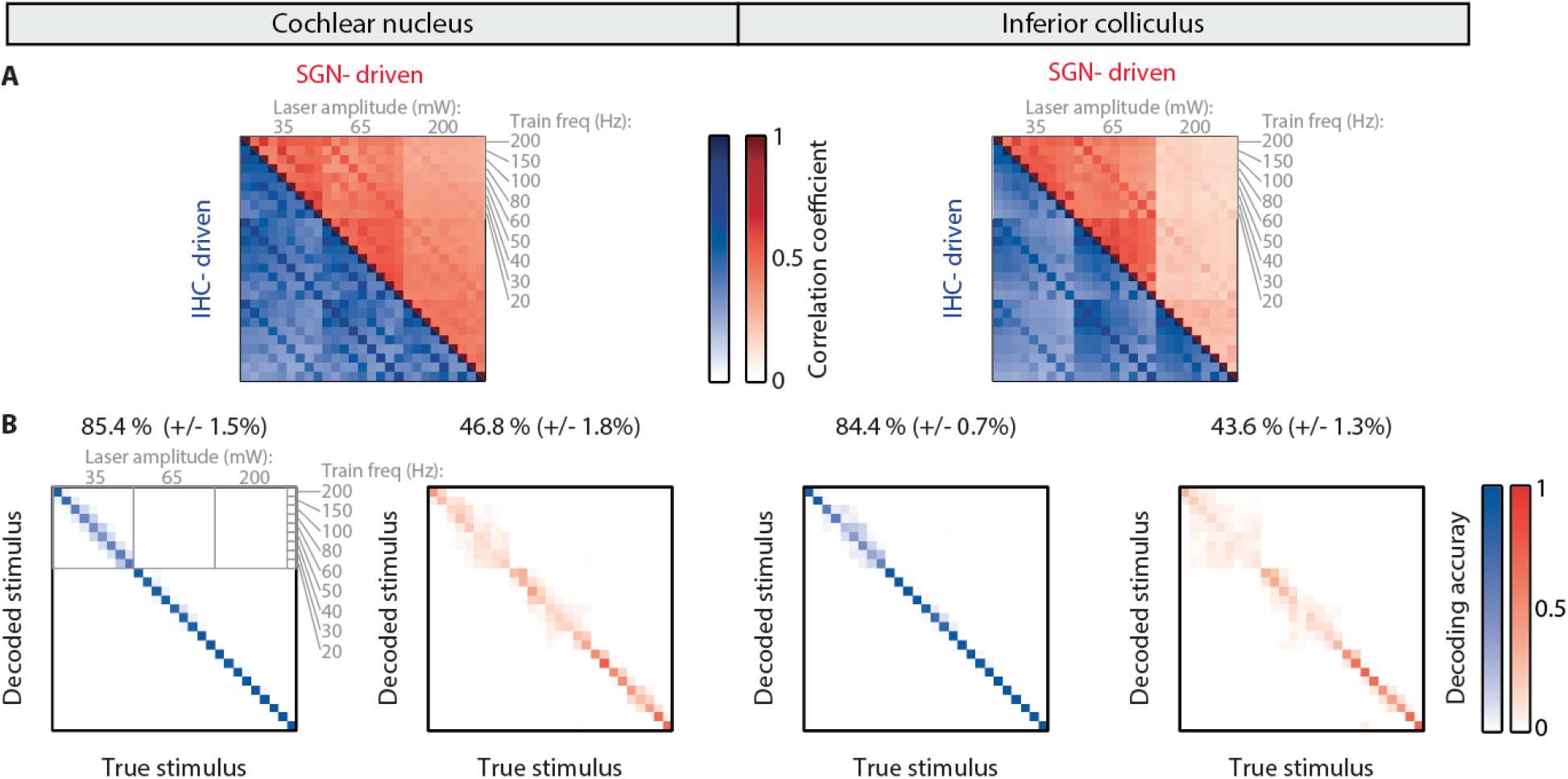
Population-level encoding of stimulation frequencies and intensities. Left half: results from the CN. Right half: results from the IC. Activation sites are denoted by color, blue: IHC-driven, red: SGN-driven (IHC stimulation: n = 630 units in CN, n = 445 units in IC; SGN stimulation: n = 335 units in CN, n = 438 units in IC). **A.** Cross-correlogram of the CN (left) and IC (right) neuronal population activity upon IHC (blue) or SGN (red) stimulation. For each condition, concatenated PSTH of all units were cross-correlated between different stimuli to obtain the similarity matrices. **B.** Confusion matrices resulting from decoding the combination of light intensity and pulse rate with a SVM classifier. We used 256 neurons in each condition out of the total neuronal pool and considered the average firing rate of each unit during the whole trial presentation for decoding. Average decoding accuracy is mentioned above each matrix (mean±STD, n = 100 random selections of 256 units, p < 10^−10^ for any comparison between IHC and SGN stimulation).

We have shown that IHC stimulation leads to better subcortical frequency tracking than SGN stimulation. Next, we asked whether this would translate into better stimulus encoding by neuronal populations. To address this question, we trained a SVM classifier to decode stimulus identity from neuronal firing rates averaged over the course of the trial’s duration (3 laser intensities × 9 pulse frequencies = 27 stimuli). Again, decoding capacity increased with the number of neurons used and plateaued after ∼200 neurons (Fig. S8A). Therefore, we fixed the population to 256 randomly selected neurons for each region and stimulation mode in subsequent analysis. The classifier performed equally well on neurons recorded from CN or IC (Fig. 5B). However, decoding accuracy was better on trials from IHC compared to SGN stimulation. In the latter case, misclassifications were frequent for trials sharing the same laser intensity. This bloc structure indicates misclassification of the modulation frequency and is consistent with what we had observed on cross-correlograms. To assess the information contained in the temporal fluctuation of neuronal discharges, we also computed the decoding accuracy from single-trial PSTH (Fig. S8B). We found that decoding accuracy was lower in every condition, especially for SGN stimulation, which suggests that firing rate rather than PSTH is more informative in this case. Finally, we turned again to channel capacity analysis and asked how many of the 27 stimuli could be faithfully recovered from subcortical population activity. We determined a channel capacity of ∼18.5 for IHC stimulation compared to ∼7.7 for SGN stimulation (Fig. S8A). We conclude that targeting IHC provided better preservation of stimulus statistics both in the intensity and frequency domains, compared to SGN stimulation.

**Figure S8:**
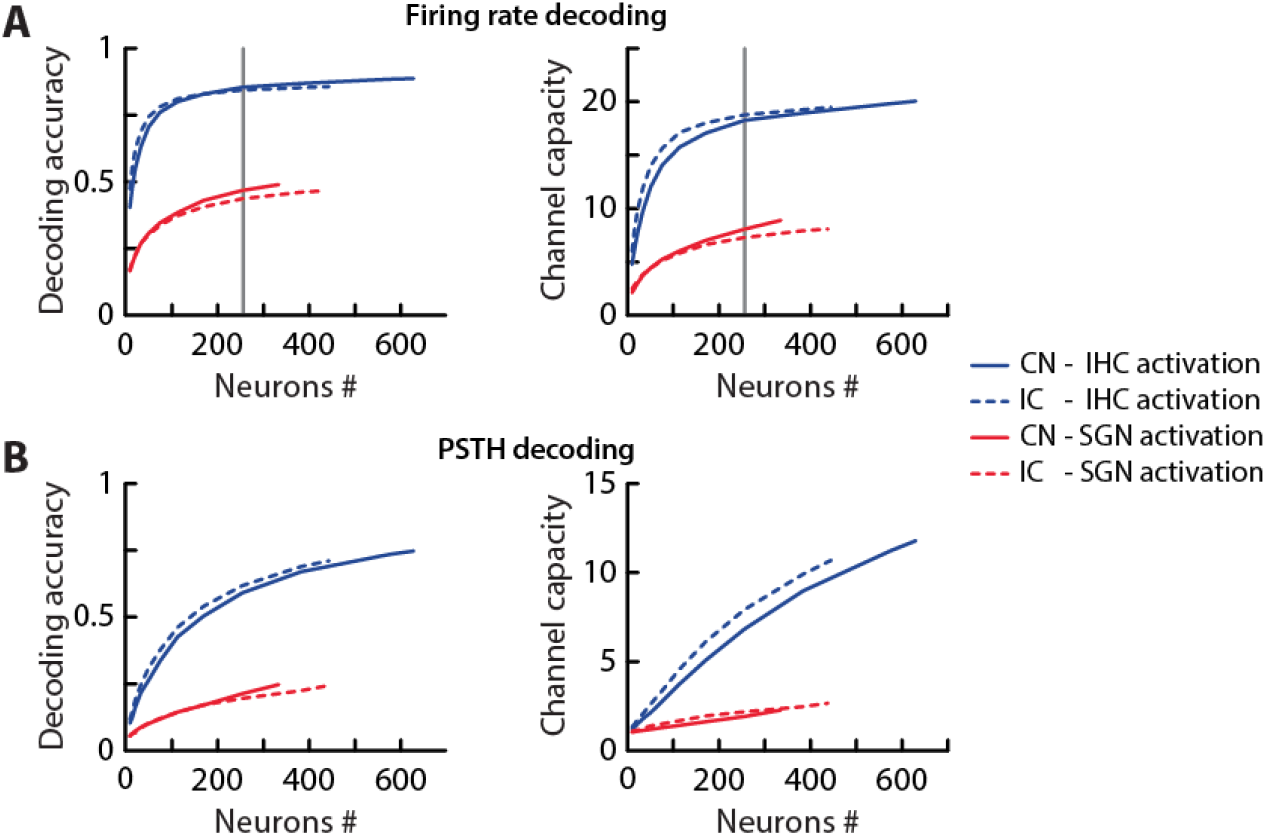
Influence of neuron numbers on the ability to decode the combination of light frequency and intensity. **A.** Decoding performance on average firing rates. The firing rate of each unit was averaged during the whole stimulus presentation before being fed to the SVM decoder. Left: Decoding accuracy of stimulus identity as a function of unit numbers for CN neurons (solid line; blue for IHC stimulation, red for SGN stimulation) and for IC neurons (dashed line; same color code). Right: corresponding channel capacity (same line and color code). Data resulted from 100 random selections of various numbers of neurons from the whole population (IHC stimulation: n = 630 units in CN, n = 445 units in IC; SGN stimulation: n = 335 units in CN, n = 438 units in IC). Note that the channel capacity is bound to N = 27 levels (3 intensity levels × 9 pulse rates). The grey line denotes a population of 256 neurons, whose confusion matrices are shown in the main text. **B.** Same as in **A** when considering temporal modulations of the firing rate. For each unit, spikes were binned over a 1 ms window to construct a single trial PSTH. Resulting time-dependent firing rates were concatenated within the population to train the decoder.

### Simulation of optogenetic activation of IHC and SGN with an auditory pathway model

To further explain our results and understand the fundamental differences between optogenetic stimulation of IHC or SGN, we turned to a computational model of the auditory pathway and adapted it to account for optogenetic stimulation (Fig. 6A). Our design was based on a previously published model that includes various stages from the spectral filtering by the basilar membrane to the firing of neurons in the brainstem (35–37). We modified several steps of this model. First, we included a biophysical description of SGN to simulate the voltage dynamics and resulting neuronal discharge. This enabled us to simulate optogenetic stimulation of these neurons using a model of channelrhodopsin kinetics. The neuronal model was based on a leaky integrate-and-fire neuron to which we added a Ca^2+^ current dynamic to account for spike-frequency adaptation. Second, we incorporated a realistic connectivity pattern between the different stages of the auditory pathway (SGN to CN and CN to IC). Third, to reproduce the experimentally observed variability of neuronal response types, we included a range of intrinsic neuronal properties and an external noise component at the different stages of processing.

**Figure 6:**
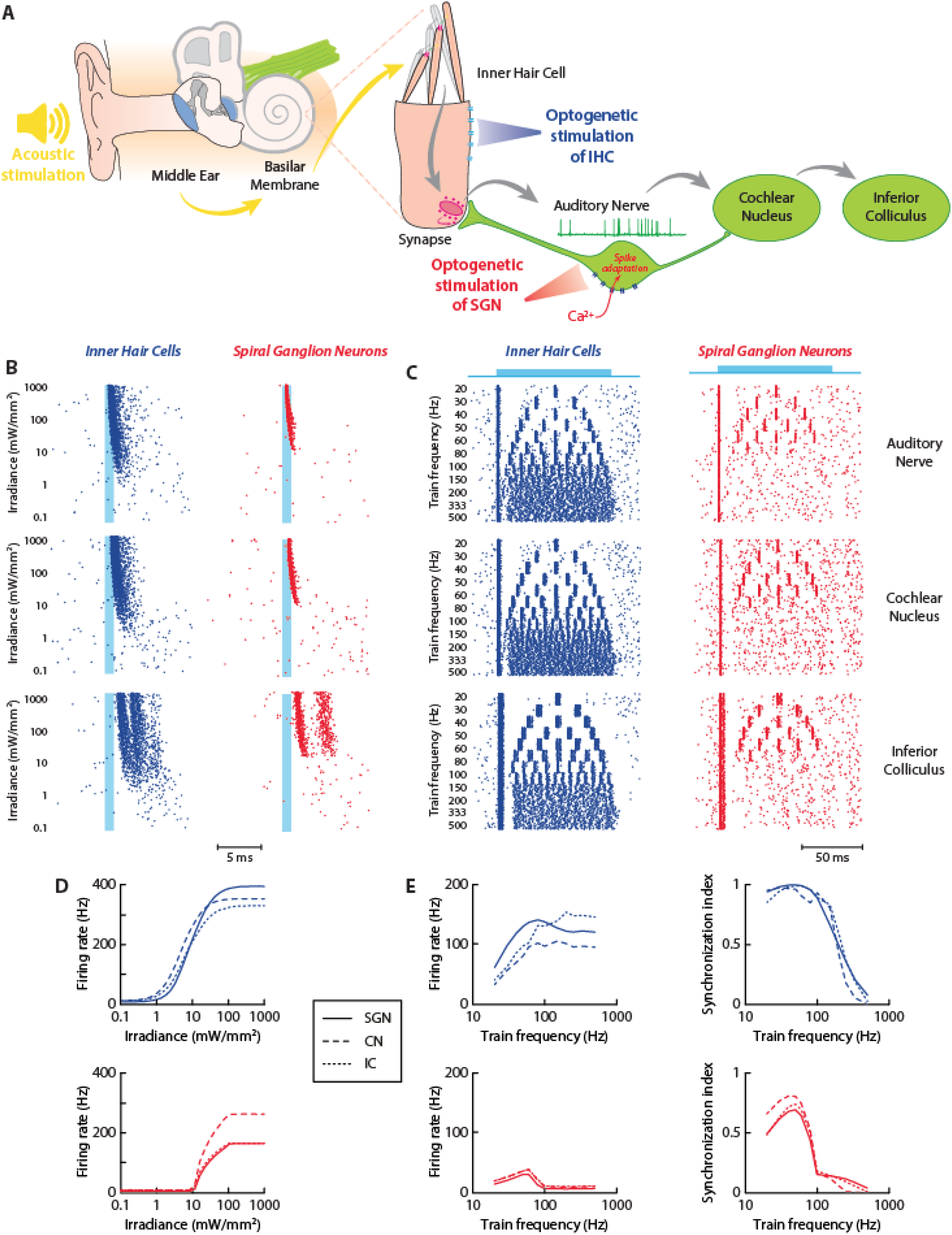
Simulation of optogenetic activation of IHC and SGN with an auditory pathway model. **A.** Schematic of the auditory pathway model. A biophysical description of spiral ganglion neurons and an optogenetic current either at the IHC or SGN sites were included. Simulations resulted in spiking activity in 3 structures: the SGN, the CN, and the IC. **B.** Spiking activity at the 3 different stages (from top to bottom) of the auditory model in response to single 1 ms light pulse of increasing power density applied to IHC (blue) or SGN (red). **C.** Spiking activity at the 3 different stages of the auditory model in response to trains of 1 ms light pulses (50 mW·mm^−2^) of increasing frequency applied to IHC (blue) or SGN (red). **D.** Firing rate in response to single 1 ms light pulses of increasing power density. **E.** Firing rate (left) and synchronization index (right) in response to trains of 1 ms light pulses (50 mW·mm^−2^) of increasing repetition rates.

The next step consisted in tuning the different parameters to reproduce basic responses to acoustic stimulations. Using pure tones of different intensities and frequencies, we verified that our network produced similar firing patterns in terms of rate and synchronization at three different stages (SGN, CN, IC). Then, we optogenetically activated either IHC or SGN (Fig. 6B-C). In response to a 1 ms light pulse, the firing rate of neurons showed a gradual increase upon IHC activation at all stages (Fig. 6D, top). For SGN activation, the response function had a rectifying shape and quickly saturated (Fig. 6D, bottom). Accordingly, the dynamic range was higher for IHC activation and decoding light intensity from the evoked firing rate was more accurate (Fig. S9A). We also found that the amount of light necessary to evoke activity was lower for IHC activation (Fig. 6D).

In response to trains of light pulses, we found fundamentally different behaviors between the two stimulation targets (Fig. 6C). For SGN activation only, the evoked firing rate showed a decrease for high repetition rates across structures (Fig. 6E, left). This was explained in our biophysical description by the fact that opsins are Ca^2+^ permeable and Ca^2+^ accumulation prevents neuronal firing past the first light pulse if the intertrial is not sufficient. We also assessed the ability to follow trains of pulses (Fig. 6E, right). Consistent with our experimental observations, we found that synchronization indices were lower after SGN compared with IHC stimulation. Accordingly, decoding was less accurate when auditory neurons were stimulated (Fig. S9B). Finally, we used our simulations to predict the effects of an opsin with faster kinetics (f-Chrimson (24, 38), Fig. S10) together with reduced Ca^2+^ permeability (Fig. S11) and found that both parameters extended the range of frequencies that the system was able to follow. Importantly, to obtain a full recovery of auditory functions in terms of dynamic range and temporal fidelity (Fig. S9), it was necessary to include both a fast temporal dynamic of the opsin and a low Ca^2+^ permeability when considering SGN stimulation.

**Figure S9:**
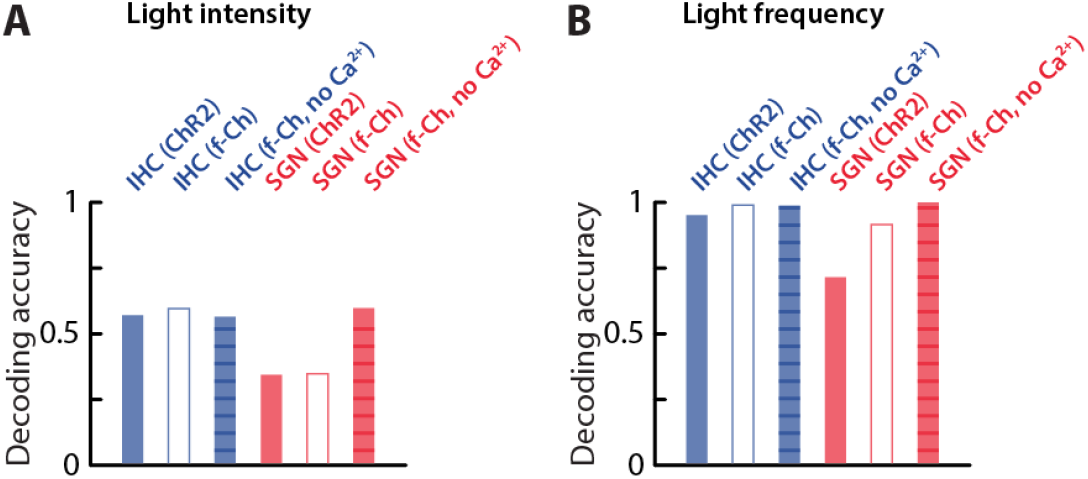
Decoding optogenetic stimulation parameters with the auditory model. **A.** Decoding accuracy of light intensity from brainstem neuron activity (decoding accuracy for CN or IC neurons was average). Light intensity took 30 logarithmically distributed values between 0.1 and 1000 mW·mm^−2^. Therefore, decoding accuracies were substantially smaller than in the experiments. **B.** Decoding accuracy of light frequency from brainstem neuron activity (decoding accuracy for CN or IC neurons was average). Light intensity took 12 different values distributed between 20 and 500 Hz.

**Figure S10:**
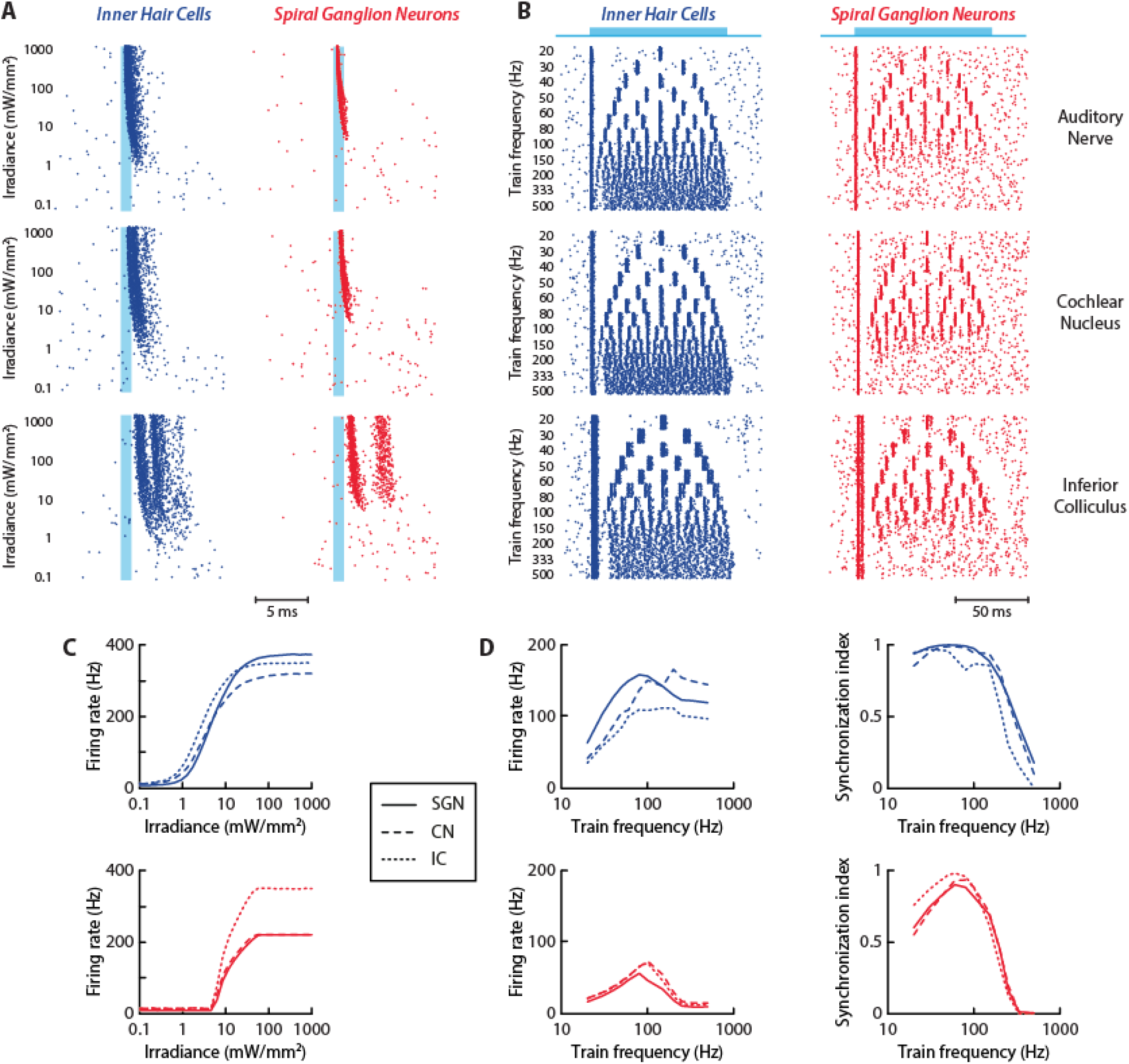
Simulation of the auditory pathway upon optogenetic stimulation with f-Chrimson. **A.** Spiking activity at the 3 different stages (from top to bottom) of the auditory model upon a 1 ms light pulse of increasing power density applied to IHC (blue) or SGN (red). **B.** Spiking activity at the 3 different stages of the auditory model in response to trains of 1 ms light pulse (50 mW·mm^−2^) of increasing frequency applied to IHC (blue) or SGN (red). **C.** Firing rate in response to 1 ms light pulses of increasing power density applied to IHC (blue) or SGN (red). **D.** Firing rate (left) and synchronization index (right) in response to trains of 1 ms light pulses (50 mW·mm^−2^) of increasing frequency applied to IHC (blue) or SGN (red).

**Figure S11:**
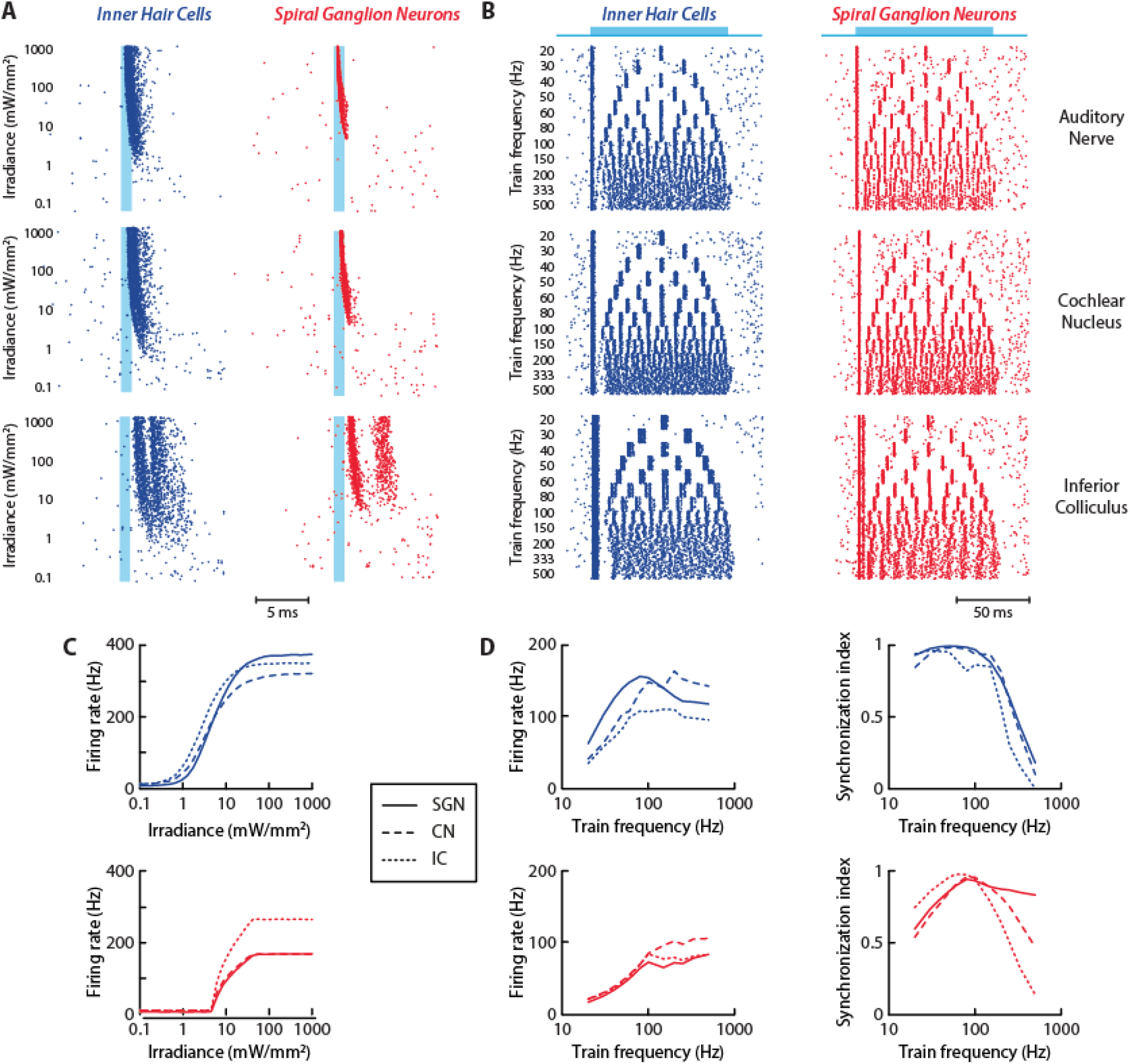
Simulation of the auditory pathway upon optogenetic stimulation with f-Chrimson and without Ca^2+^ current. **A.** Spiking activity at the 3 different stages (from top to bottom) of the auditory model upon a 1 ms light pulse of increasing power density applied to IHC (blue) or SGN (red). **B.** Spiking activity at the 3 different stages of the auditory model in response to trains of 1 ms light pulse (50 mW·mm^−2^) of increasing frequency applied to IHC (blue) or SGN (red). **C.** Firing rate in response to 1 ms light pulses of increasing power density applied to IHC (blue) or SGN (red). **D.** Firing rate (left) and synchronization index (right) in response to trains of 1 ms light pulses (50 mW·mm^−2^) of increasing frequency applied to IHC (blue) or SGN (red).

To comprehensively validate the model and our experimental strategy, more complex auditory stimuli were required. The ESC-50 environmental sound dataset, which contains 2,000 audio recordings across 50 categories (including animals, natural sounds, human non-speech sounds, household noises, and urban sounds), was used to evaluate the model under different stimulation strategies and channelrhodopsin variants (Fig. 7). To simulate optogenetic stimulation, acoustic signals were first converted into cochlear implant-like stimulations using a filter bank and envelope extraction. Because continuous stimulation caused channelrhodopsin to remain in a light-adapted state and lose responsiveness, minimum and maximum stimulation thresholds were introduced to allow recovery periods and maintain effective activation. Finally, neural responses from the SGN stage (neurogram, Fig. S12) were used to train a Convolutional Neural Network (CNN) decoder and classification accuracy was assessed on the whole dataset.

**Figure 7:**
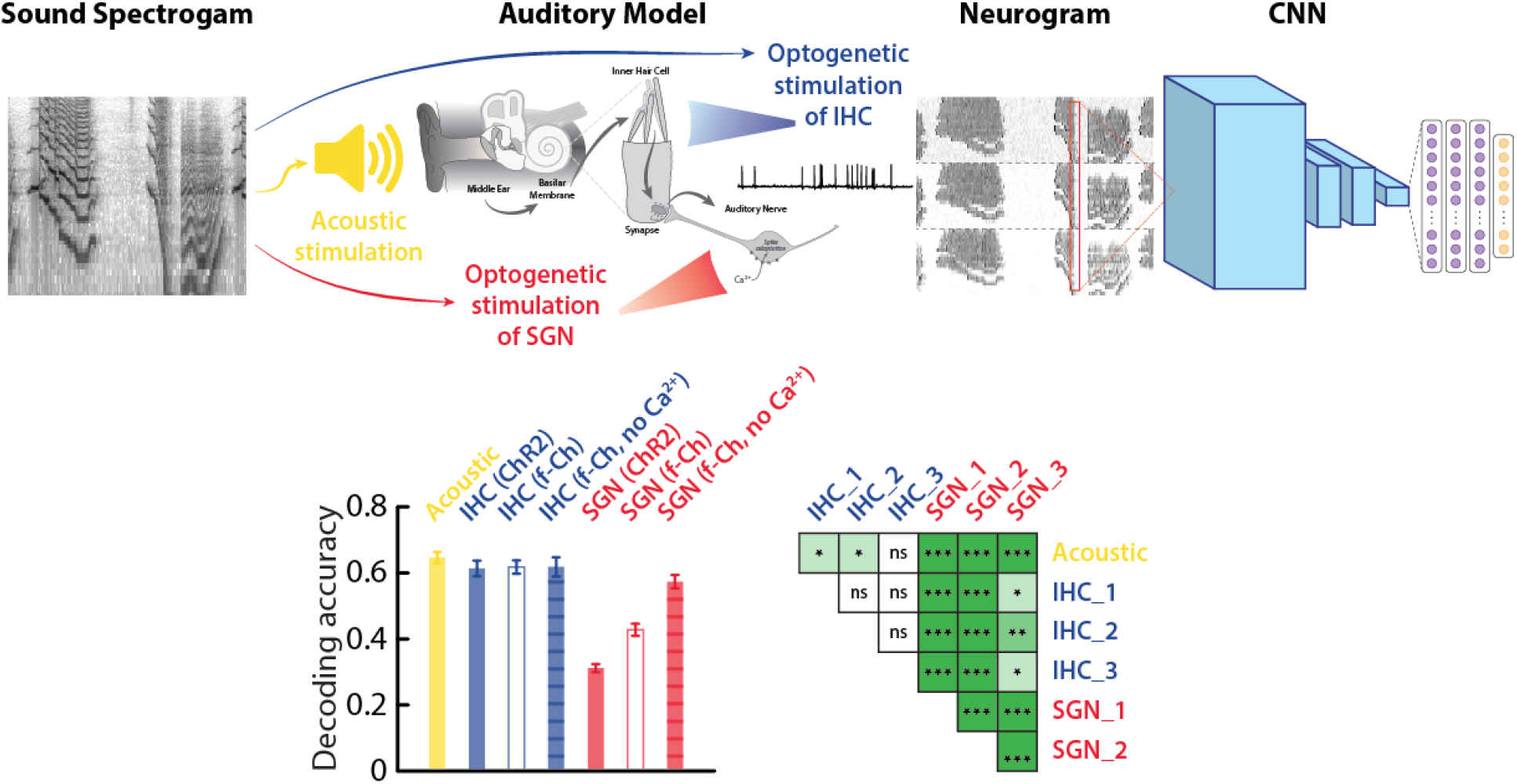
Response of the model to natural sounds and decoding with a convolutional neural network. Top, Schematic of the simulation and decoding workflow: the 2,000 sound waveforms were processed by different auditory models (acoustic, yellow; optogenetic activation of IHC, blue; optogenetic activation of SGN, red). The resulting neurograms were binned and used as inputs to the Convolutional Neural Network to decode stimuli identity. Bottom, decoding accuracy (mean±STD) for the different models and corresponding statistical significance of all comparisons (T-test statistics, number of observations is the number of folds in the decoding step: n = 5).

The acoustic reference model achieved 64.6% accuracy (Fig. 7) which is identical to what could be decoded directly from the audio spectrogram with a similar previously developed CNN (39). Optogenetic stimulation of IHC produced very similar performance (≈ 61-62%) regardless of the channelrhodopsin variant, indicating that this approach preserves the temporal and spectral information required for accurate sound encoding. In contrast, SGN stimulation resulted in lower and more variable performance, strongly depending on the opsin used: ChR2 reached 31.2%, f-Chrimson 42.8%, and f-Chrimson without Ca^2+^ current 57.3%. These results demonstrate that stimulating sensory hair cells is consistently more effective than direct auditory nerve stimulation. It underlies the importance of natural synaptic processing for preserving auditory information, while also showing that optimized opsins can substantially improve auditory nerve optogenetic stimulation.

**Figure S12.**
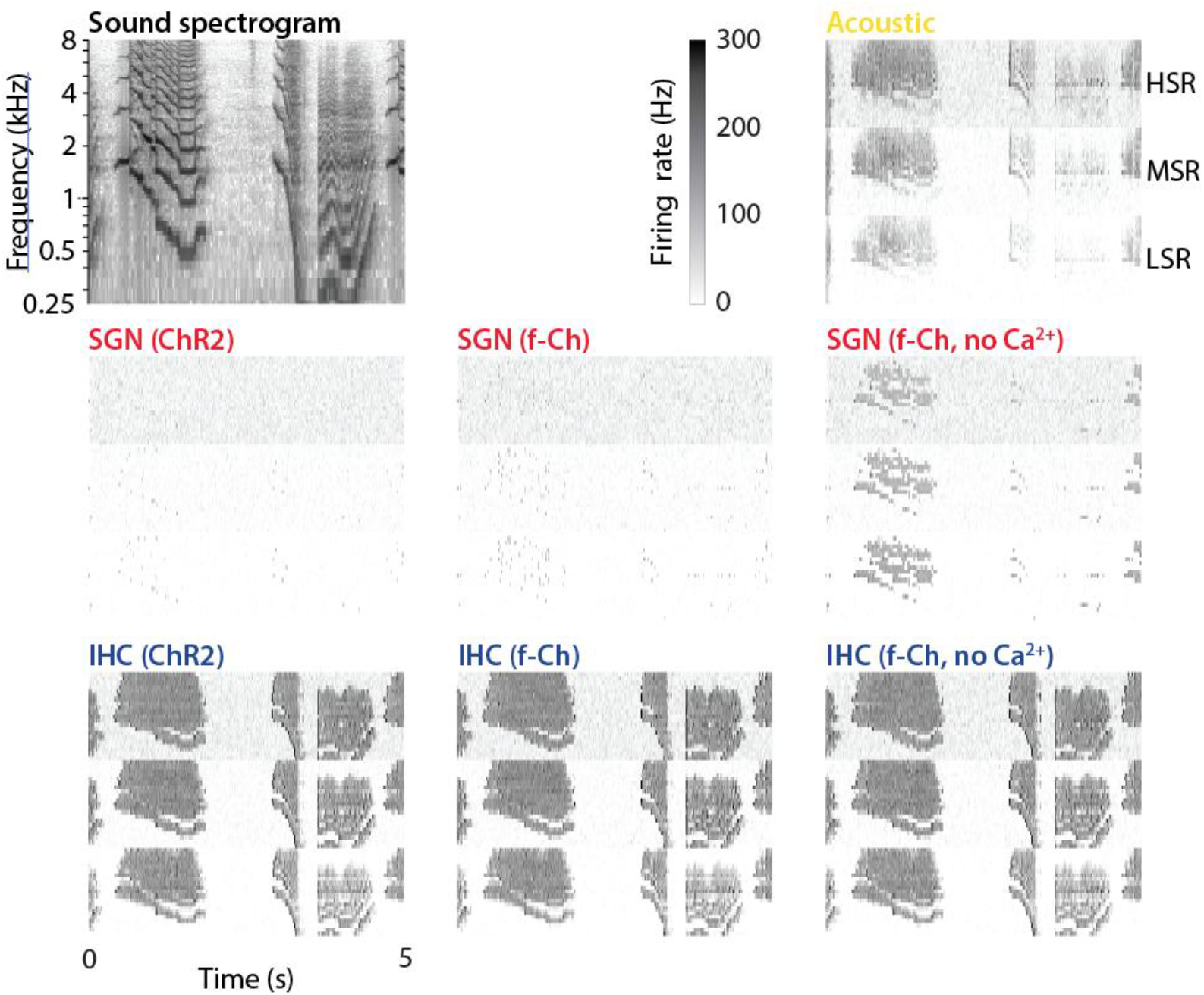
Neuronal activity in the different configurations of the auditory model. Processed neurograms used as input to the CNN. Neuronal activity was simulated by the 7 different models (acoustic, SGN stimulation, and IHC stimulation with the different opsin variants) in response to a sound stimulus, whose spectrogram is represented on the top left. Neurons are organized according to their spontaneous firing activity (HSR, MSR, LSR for high, medium, and low spontaneous rates).

## DISCUSSION

Electrical cochlear implants (eCI) have enabled over one million patients (1) suffering from profound hearing impairment or deafness to recover some degree of speech comprehension and significantly improve their quality of life. Yet, the hearing restoration offered by eCI is rudimentary owing to limitations in spectral and intensity coding. Optical stimulation of SGN has been brought forward as a promising alternative to eCI, as it offers superior spatial confinement of stimulation. Here, we build on this concept and extend it to optical stimulation of sensory IHC. In mice, we compare the two targets by recording optically evoked ABR and multi-unit neuronal activity in two brainstem relays. We demonstrate that compared to SGN optogenetic stimulation of IHC better preserves the temporal and intensity content of stimuli across processing stages.

### Relation to previous work on cochlear activation via optogenetics

Initial approaches for optogenetic activation of the cochlea have used several different genetic strategies. Initially, the generic mouse line expressing ChR2 in a large number of neuron types under the *Thy1.2* promoter (40) has been used to assess the possibility of optogenetic stimulation of the auditory nerve (10). Transgenic animals driving ChR2 expression under the *parvalbumin* promoter showed reliable expression both in IHC and SGN and good entrainment to pulse train stimuli were observed in auditory nerve or IC neurons (41, 42). For selective expression in IHC, the *Vglut3* promoter has been used to characterize light-induced synaptic fusion at the ribbon synapse *in vitro* (28). However, ectopic expression in SGN has also been reported, in agreement with our present results (Fig. S2). Although selectively targeting ChR2 to OHC is achievable with the *prestin* promoter, it failed to elicit physiologically relevant effects (25), potentially because the dense membrane expression of *prestin* may compete with ChR2 membrane trafficking (43). Lastly, ChR2 has been expressed in SGN with the *Bhlhb5* promoter, but mostly for characterizing the electrophysiological properties of these neurons *in vitro* (44). The diversity of transgenic approaches that have been explored highlights that finding promoters to specific cell type remains challenging, especially for auditory neurons.

Next to transgenic models, viral delivery of opsins via cochlear injection of AAV vectors represents an interesting alternative as it offers a certain degree of spatial confinement and is more readily translatable to clinical settings. Several studies have reported successful viral targeting of opsins to SGN using the *hSyn* promoter. However, off-target expression is often observed in IHC of the gerbil model (45, 46), thereby confounding the interpretation of data. Nonetheless, one important advantage of AAV-driven expression is the versatility of opsins that may be expressed. The CatCh variant with increased photocurrent and higher Ca^2+^ permeability has been employed in several studies (10, 12, 45, 46) but remains relatively slow for auditory applications. Given our simulation results, we would suggest that a less Ca^2+^ permeable opsin might be more appropriate for SGN stimulation. Red-shifted variants such as ChRmine or ChReef are promising for substantially decreasing the light threshold and shifting the excitation spectrum toward the red but at the cost of lower temporal fidelity (47). Considerable efforts have been made to develop opsin variants with faster timescale and red-shifted excitation spectrum and the family of f-Chrimson variants have emerged as optimal tools for auditory applications (24, 38). Results showed the ability to follow temporal patterns up to ∼200 Hz similar to IHC stimulation with a slower opsin in our study. Therefore, it would be interesting to express this variant in IHC and determine the potential benefits.

Here, we aimed to faithfully characterize and compare the contribution of optically driven SGN or IHC inputs to the fidelity of stimulus-encoding. Because off-target expression in some studies precludes any definitive conclusion, we carefully chose Cre-driver lines, *Bhlhb5^Cre^* and *Myo15a^Cre^*, that enabled exclusive expression of ChR2 in either of the two targeted populations (29, 30) and used identical protocols to compare their activation properties.

### Optogenetic activation of inner hair cells

With this approach, we show that optogenetic IHC stimulation robustly activates the auditory pathway up to the midbrain. This may be explained by the signal amplification at the IHC-to-SGN synapse, where a single IHC maintains contact with 10-30 SGN, each representing an individual synaptic connection. Our results also demonstrate the advantages of IHC compared to SGN stimulation, in terms of sensitivity, dynamic range (Fig. 3) and temporal fidelity (Fig. 4). In experiments, the decrease in threshold may be explained by several factors, such as a higher opsins density or the closer proximity of IHC to the optic fiber. Direct quantitative comparisons with previous studies or with simulation results remain difficult because optical thresholds depend on the surgical approach, light delivery geometry, opsin expression, and the biophysical properties of the opsins used. Notably, our experiments were performed without cochleostomy, suggesting that substantially lower thresholds could be achieved by placing the light source directly within the scala media. In addition, an intrinsic cellular mechanism may contribute to the enhanced sensitivity. Ca^2+^ entering through channelrhodopsin could directly trigger synaptic vesicle release, partially bypassing the need for large membrane depolarization and activation of endogenous voltage-gated Ca^2+^ channels. Such a mechanism could contribute to the expanded dynamic range observed with IHC stimulation. A recent report examined in a systematic manner the dynamical range reached by different opsins and found values around 13 dB (48), a value very similar to what we found after SGN activation. Here, we demonstrate that the dynamic range could be increased by 50% through IHC activation.

It is noteworthy that direct stimulation of sensory hair cells is a closer match to physiological conditions as many features of synaptic transmission and natural spiking of SGN are conserved. In particular, reliability, dynamics, and adaptation of synaptic transmission, which are central for proper synaptic signaling and underlying processing (49, 50), were preserved with our optogenetic approach. Furthermore, IHC are contacted by different classes of afferent fibers with distinct sensitivities to sound intensity (51, 52). This mechanism that permits to maintain the large dynamic range of hearing is lost when SGN are activated directly, thereby limiting intensity coding. Accordingly, although optogenetic activation of SGN achieves better spatial resolution than eCIs (8), it is still far from matching the performance of natural hearing and additional improvements may be reached through the direct activation of sensory hair cells.

Some of the limitations related to direct activation of SGN may be improved by exploiting next-generation opsins with faster kinetics (38, 53). Recent studies using f-Chrimson expressed in SGN of gerbils reported near-physiological encoding efficiency of temporal stimulus statistics in the IC (48). Interestingly, this is in line with the predictions obtained by simulating f-Chrimson within our auditory pathway computational model (Fig. 6-7). However, our simulations indicate that improvements in opsin kinetics alone are unlikely to fully compensate for bypassing the IHC-SGN synapses. Additionally, efficiency of optogenetic stimulation is often assessed using simple and tractable stimuli such as rectangular light pulses of various intensity and repetition rates. We showed in our simulations that although decoding accuracy of these variables may be equally good after IHC or SGN stimulation with optimized opsins (Fig. S9), SGN stimulation did not reach the same level of performance when considering a more complex dataset of natural sound stimuli (Fig. 7).

### Application for hearing restoration

Direct stimulation of SGN as a proof of principle for future oCI (10, 11, 16) has the advantage of covering a larger spectrum of etiologies underlying hearing loss, including those that leave patients without functional IHC. Indeed, death or degeneration of IHC and/or peripheral afferents are common during deafness and hearing loss in both humans (54, 55), and mice (56, 57). However, not all forms of hearing loss involve complete loss of IHC. During age-related hearing loss, degeneration is generally more pronounced in OHC than in IHC (56, 58) and several congenital forms of hearing loss, including Usher syndromes 1 and 2, affect the mechano-transduction process but could leave the hair cell soma intact as a potential target for therapeutic intervention (59).

The feasibility of these two strategies also differs from a gene delivery perspective. Efficient optogenetic transduction of adult SGN using AAV remains technically challenging and has so far been achieved reliably only in gerbils, but not in mice (6). In contrast, numerous AAV capsids and promoters have recently been developed and enable efficient and selective transduction of IHC, providing a broader toolbox for therapeutic applications (60).

Our results also support the feasibility of targeting IHC in the context of hearing loss. Using C57BL/6J mice, which carry a mutation in the *Cdh23* gene causing progressive hearing loss (61), we observed robust optically evoked auditory brainstem responses in animals up to 70 weeks of age. This demonstrates that the IHC-SGN synapses are preserved in this genetic model of auditory ageing. Thus, SGN and IHC target-based approaches for the development of future oCI may complement each other to cater for the needs of distinct patient groups, while optimizing performance and efficiency.

Finally, targeting IHC may simplify the implant design. Targeting light efficiently to IHC or SGN is challenging due to the cochlea’s intricate anatomy. Because IHC would be closer to the implant than SGN, this would reduce the necessary light intensity and limit energy consumption as well as heat production. Indeed, our results show that IHC activation requires less light power and comes to support IHC as a promising target for the development of more efficient and safer implants with a lower energy footprint and superior longevity.

### Limitations and future perspectives

Although our data strongly supports IHC as a promising target for the development of more performant oCI, this remains to be engineered and tested pre-clinically. A considerable challenge when targeting IHC is to preserve sensory hair cell integrity during the implantation process. Indeed, implanted human patients often retain some residual hearing, but mostly at locations apical to the tip of the electrode (62). Nevertheless, recent pharmacological developments promoting protective effects on hair cells have demonstrated a substantial survival of IHC and afferent synapses in animal models (63). Technological advancements have been pivotal in the development of oCI. Researchers have designed high-density micro-light-emitting diode (μLED)-based oCI with optimized thermomechanical properties suitable for optogenetic experiments. These devices comprise numerous miniaturized LED distributed along a flexible probe, allowing for precise spatial activation of the auditory nerve fibers (64). Further improvements in this direction may be useful to decrease the spatial footprint of the implant and preserve sensory IHC for optical stimulation.

Additional challenges include the long-term safety of optogenetic approaches, particularly regarding potential toxicity and immune responses associated with viral delivery of opsins. These concerns may be especially relevant for IHC-targeted strategies, as these highly specialized sensory cells must maintain their structural integrity and synaptic function over long periods. Continued development of safer viral vectors, improved promoters, and more selective gene delivery strategies will therefore be essential.

Nevertheless, recent advances in gene therapy and its successful use for hearing (65) or vision (66) restoration, highlight the growing potential of optogenetic strategies for treating sensory disorders. Optogenetic implants offer the potential to enhance hearing perception through more accurate activation of tonotopic regions of the cochlea (7, 8). Here, optically targeting IHC rather than SGN, we further augment the performance that can be achieved in terms of dynamic range and temporal resolution. We conclude that an oCI targeting IHC would considerably increase the amount of information that can be conveyed and would expect improvements for speech comprehension, hearing in noisy environments, and listening to complex sounds such as music.

## MATERIALS AND METHODS

### Animals

All experimental procedures were performed in accordance with French and European regulations for the care and protection of laboratory animals (EC Directive 2010/63, French Law 2013–118, February 6, 2013), under authorization from the Institut Pasteur’s Ethics Committee for Animal Experimentation. The use of transgenic animals was in agreement with the European directive 2009/41/EC (French Law 2013-1177 and 2021-1905). The animals were placed in standard housing in 12h light/12h dark conditions, with a background noise < 40 dB SPL. They had unlimited access to food and water and were housed in cages with a maximum of five mice each.

For expression of ChR2(H134R)-tdTomato (67), we used the Cre-loxP system for optogenetics (68). We crossbred the B6.Cg-*Gt(ROSA)26Sor^tm27.1(CAG-COP4*H134R/tdTomato)Hze^*/J mouse line (abbreviated as Ai27D; IMSR_JAX:012567), with three different Cre-expressing mouse lines. To express the opsin in SGN, the Cre recombinase was under the control of the *Bhlhe22* promoter using the *Bhlhe22^tm2(cre)Gan^*mouse line (abbreviated as *Bhlhb5^Cre^*; MGI:3830517; ref. (30)). To express the opsin in IHC, the Cre recombinase was under the control of the *Myo15a* promoter using the *Myo15a^tm1.1(cre)Ugds^*mouse line (abbreviated as *Myo15^Cre^*; MGI:4361284; ref. (29)). To express the opsin in both IHC and SGN, the Cre recombinase was under the control of the *Vglut3* promoter using the B6;*129S-Slc17a8^tm1.1(cre)Hze^/J* (abbreviated as *Vglut3^Cre^*; MGI:5792821). The present study made use of 42 animals for *in vivo* experiments. Recording of optogenetic ABRs used animals within 6 to 70 weeks of age (number of animals for the different strains: *Bhlhb5^cre^ n = 4, Vglut3^cre^ n = 15, Pmyo15^cre^ n = 23)*. Intracranial recordings were exclusively done on young animals aged from 6 to 12 weeks (number of animals for the different strains: *Bhlhb5^cre^ n = 5, Pmyo15^cre^ n = 6*). For *in vitro* experiments (patch-clamp on hair cells), we crossbred the Ai27D mouse line with the B6.Cg-Tg(Atoh1-cre)1Bfri/J mouse line (abbreviated as *Atoh1^Cre^*; MGI:J:102293) to express the opsin both in inner and outer hair cells. Experiments were performed on an excised preparation of the organ of Corti using animals aged between P10 and P13.

Males and females were included. No inclusion/exclusion criteria were applied, and no outlier removal was performed. Animals were randomly selected for experiments. Sample numbers and experimental replicates for all studies are provided in the figure legends.

### *In vitro* electrophysiology

Recordings were performed at room temperature in HEPES-based extracellular solution that contained (in mM): 143 NaCl, 5 D-glucose, 6 KCl, 1.3 CaCl_2_, 2 Na-pyruvate, 0.7 NaH_2_PO_4_, 0.9 MgCl_2_, and 10 HEPES. Electrodes, pulled from borosilicate pipettes (1.5 OD) on a Flaming/Brown micropipette puller (Sutter Instruments), had resistances in the range of 6-10 MΩ when filled with internal solution containing (in mM): 130 KCl, 10 NaCl, 3.5 MgCl_2_, 1 EGTA, 5 ATP-K_2_, 0.5 GTP-Na_2_., and 5 HEPES.

Cells were visualized through a ×20 water-immersion objective using infrared differential interference contrast and fluorescence microscopy (BX51, Olympus). Whole-cell current-clamp recordings were made using BVC-700A amplifiers (Dagan). The signal was filtered at 5 kHz and digitized at 25 kHz using an 18-bits interface card (PCI-6289, National Instrument). In order to activate only the cell of interest, optical stimulation was performed using a custom made pattern generator (69) based on a Digital Micromirror Device (DLP LightCrafter; Texas Instrument). We used the inbuilt blue LED of the projector which has a center wavelength of 460 nm and intensity of 10 mW·mm^−2^ at the sample plane. Signal generation and acquisition were controlled by a custom user interface programmed with LabVIEW (National Instrument).

### Acoustic and optical stimulations

Acoustic stimuli were generated via a high-frequency auditory signal processor (RZ6, Tucker-Davis Technologies). The processor was used for generating acoustic stimuli via a multi-field magnetic speaker (MF1, Tucker-Davis Technologies) and triggers for laser stimulation (Cobolt 06-01 473 nm, Hübner Photonics). An optical fiber of diameter 100 µm (M63L01, Thorlabs) was cleaved and polished to make a flat-cleaved end before being used for photostimulation. The maximum light intensity was measured with a photodiode power meter (S12C, Thorlabs) and took a value of 200 mW. Signal generation and visualization were operated via a custom graphical user interface developed in LabVIEW (National Instruments).

### *In vivo* electrophysiological recordings

Surgery and recordings were performed under general anesthesia (1.5-2% isoflurane at a flow rate of 0.2 L·min^−1^ of 95% O_2_) and analgesia achieved through local application of xylocaine (5 mg·kg^−1^) and intraperitoneal injection of buprenorphine (0.1 mg·kg^−1^, 30 min before surgery). The animal was placed on a stereotactic mask in an acoustically and electromagnetically isolated chamber.

Preceding each recording, the animal’s hearing thresholds were measured with ABR. The subcutaneous electrodes were connected to a Medusa4Z preamplifier (Tucker-Davis Technologies). The RZ6 also transmitted the sounds to a free-field (TDT MF1) transducer. Signals were filtered between 3 Hz and 3 kHz and further processed using a custom-built graphical interface (LabVIEW, National Instruments).

To gain visual access to the cochlea, we ablated the pinna, opened the ear canal, removed the eardrum, malleus and incus, and enlarged the bulla opening using a micro Friedman-Pearson rongeur (Fine Science Tools). The optical fiber was approached to the middle turn of the cochlea without any cochleostomy and optical ABR were monitored.

After measuring optical ABR, we performed a craniotomy above the contralateral IC. A Neuropixels 1.0 probe (IMEC) was inserted at a location defined by −5.02 mm AP, 1.72 mm ML with a 43° elevation angle from horizontal and a 95° azimuth angle from midline. After piercing the dura, the probe was lowered at a speed of 20 µm·s^−1^ traveling through the contralateral IC and until reaching the ipsilateral CN (∼5 mm). Neuronal responses to short (1 ms) light pulses of moderate intensity (20 mW) were monitored throughout the insertion process to check the correct placement of the electrode.

Once the electrode was in place, we presented 1) single light pulses of 1 ms at 13 different light intensities (from 0.2 mW to 200 mW, logarithmically spaced) and 2) 100 ms pulse trains at 9 different rates (1 ms single pulse; 20, 30, 40, 50, 60, 80, 100, 150, 200 Hz) and 3 light intensities (35, 65, 200 mW). Recording sessions consisted in randomly interleaved repetitions of 50 trials of each stimulus separated by 800 ms intertrial intervals to allow recovery of the opsin. Recordings lasted ∼30 min after which we retracted the electrode and implanted it again at a nearby location. We were able to perform 1-3 recordings per animal.

To verify the correct placement of the electrode, the probe was coated with DiI dye after the last recording and reinserted into the same location for 20 minutes. Subsequently, standard histology procedures were performed. The brain was dissected and post-fixed in 4% paraformaldehyde (PFA) overnight, washed four times in phosphate-buffered saline (PBS) and cryoprotected in a 30% sucrose solution. Tissues were frozen in isopentane at −26 °C and cryosectioned at 35 µm. Brain sections were counterstained with DAPI and imaged on a Zeiss confocal microscope (LSM 900 Airyscan). Image series were registered to the Allen CCF Atlas using a Matlab registration tool (AP_histology by Andy Peters, https://github.com/petersaj/AP_histology) to confirm the probe placement within the targeted auditory structures.

### Data processing

Data preprocessing involved spike sorting using Kilosort 3, followed by manual curation with Phy2 (70). Units either exhibiting modulated post-stimulus time histogram (PSTH) or increased firing rate by 3 standard deviations above the pre-trial period were considered responsive. For all recordings, responsive units were consistently located within the range of the CN or IC, determined by stereotaxic coordinates and verified by histology in 2 animals of each genotype.

### Data analysis for single pulse stimulations

#### Onset threshold and dynamic range

For each unit in the CN or IC, the average firing rate was measured between 1.5 and 25 ms post stimulus onset. The onset threshold was defined as the necessary light intensity to evoke a firing rate exceeding the baseline by 3 STD. The dynamic range was defined by response function width from 10% above the baseline to 90% of the maximum firing rate. The light intensity in dB was defined as *I_dB_* = 20 · log(*I_W_*⁄*I*_0,*W*_), where *I_W_* is the light power and *I*_0,*W*_ ≈ 200 mW is the maximal achievable power at the optic fiber tip with our laser source.

#### Decay time constant

Data were binned with a 1 ms window to obtain the trial averaged PSTH. We then computed the autocorrelation of the PSTH across 100 ms time lags before fitting a decaying exponential function. We defined the neural timescale (*τ*) for each unit as the decay time constant derived from the fit. For units where a single-exponential fit accounts for less than 75% of the variance, we applied a double-exponential fit *f*(*t*) = *a*_1_*e*^−*t*/*τ*1^ + *a*_2_*e*^−*t*/*τ*2^, and calculated the weighted sum of the two constants as: *τ* = (*a*_1_ · *τ*_1_ + *a*_2_ · *τ*_2_)⁄(*a*_1_ + *a*_2_). This approach aligns with previous research (71). All fits were visually inspected for accuracy.

#### Decoding

We evaluated the ability of the neuronal population to decode the stimulus identity using a Support Vector Machine linear classifier. Single trial firing rate for each unit was first computed within the same 1.5-25 ms measurement window. We trained a linear-kernel decoder in a one-vs-rest scheme and five-fold cross-validation on a fraction ([10, 15, 22, 34, 51, 76, 114, 171, 256, 384, 576] neurons) of the data or on all data. The process was repeated 100 times on a random subset of neurons to compute and average decoding accuracy and confusion matrix. The average confusion matrix was used to estimate the channel capacity using the Blahut-Arimoto algorithm.

### Data analysis for pulse train stimulations

#### Synchronization index

For each unit at every laser train rate and intensity, we computed the synchronization index (*SI*) to quantify the degree of phase-locking. To avoid bias from the onset transient response, we excluded all spikes occurring within the first 25 ms after stimulus onset. *SI* was calculated as:

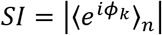

where *φ_k_* is the phase of spike *k* relative to the modulation cycle.

The statistical significance of this measure can be estimated from the Rayleigh statistic *R*:

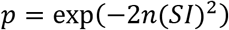

We considered synchronization to be statistically significant if *p* ≤ 0.01 which corresponds to *SI* ≥ 0.21 with *n* = 50 trials (72).

### Principal Components Analysis

Units from both groups were pooled. For each of the 27 stimulus classes (9 laser rates × 3 intensities), spike times were binned at 1 ms and averaged across trials to obtain a PSTH. The 27 PSTH were concatenated and stacked across units to form a matrix of shape: rows = classes × time bins and columns = units. Each column (unit) was Z-scored across all rows. PCA was applied to this matrix, and the number of retained components was chosen by visual inspection of the scree “elbow.” For visualization, PC scores were reshaped to laser rate × intensity × time bin × number of PC. To quantify group contributions, we summed the absolute values of each unit’s loading of a given group and divided by the sum across all units.

#### Cross-Correlogram

To assess the similarity of neural population responses across stimulus conditions, we considered the 27 stimulus classes, as for the PCA analyses. For each unit and stimulus condition, we first computed the Z-scored PSTH of each unit by subtracting its baseline mean and then dividing by the baseline standard deviation. PSTH from all units of a given brain region and for a given stimulation target were concatenated and pairwise correlations were then calculated between these combined PSTH.

#### Decoding

Single trial firing rate (average with the 0-120 ms measurement window) or PSTH (binned over a 1 ms window) for each unit were used to train a SVM linear classifier. Resulting scalars or vectors were concatenated across units. As in the previous section, a fraction ([10, 15, 22, 34, 51, 76, 114, 171, 256, 384, 576] neurons) of the data or all data was used for decoding and channel capacity estimation.

### Computational model

To interpret the optogenetically-evoked responses recorded in the cochlear nucleus and inferior colliculus, we used a customized version of an auditory periphery and brainstem model (35–37). The model includes middle-ear filtering, basilar-membrane mechanics, IHC receptor potentials, stochastic ribbon-synapse, SGN fibers, and chopper-type neurons in CN and IC (Fig. 6).

#### Leaky-integrate-and-fire neuron model of spiral ganglion neurons

In the original model, neuronal firing in the spiral ganglion neurons happened at each synaptic fusion at the presynaptic site, modulo a refractory period. In order to simulate optogenetic activation of these neurons, we replaced this simplified process with a biophysical description of the neurons. For this purpose, we used a leaky-integrate-and-fire (LIF) description of neuronal dynamics. We included 4 contributions of different currents likely to affect the membrane potential:

- An excitatory postsynaptic current *I_syn_* resulting from vesicle release. Excitatory postsynaptic conductance produced by a vesicle fusion was converted to an injected current using a single-exponential convolution function of amplitude 0.5 nA and decay τ = 1 ms (73).
- A photocurrent *I_opto_* = *g_C_*_ℎ*R*2_ · *V_drive_* where *g_C_*_ℎ*R*2_ is generated by a four-state ChR2 model with inward rectification and *V_drive_* is the driving force of the optogenetic current (with parameters identical as in the publication). For the original ChR2, we left all Markov-model rate constants unchanged, and hand tuned only the maximal conductance *g_C_*_ℎ*R*2,*max*_ to our own patch-clamp recordings of light evoked current in IHC (Fig. S1). For ultrafast red-shifted variants (f-Chrimson), both the transition rates and the maximal conductance were adjusted to reproduced published kinetics and amplitudes of optogenetic activation (38, 74). We implemented three channelrhodopsin variants that had different kinetics and Ca^2+^ permeabilities:

- wild-type ChR2 (5% Ca²⁺ permeability),
- f-Chrimson, a red-shifted ultrafast opsin (2.5% Ca²⁺ permeability),
- a hypothetical Ca²⁺-impermeable mutant (0% Ca²⁺ permeability).
- A spike adaptation mechanism based on a Ca^2+^-dependent afterhyperpolarization current *I_AHP_* (75). Intracellular Ca^2+^ concentration, [Ca], was governed by first-order kinetics according to:

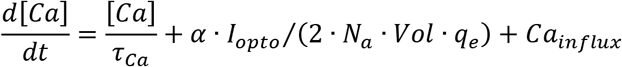

with the calcium-clearance time constant *τ_ca_* set to 5 ms. *N_a_* is the Avogadro number, *q_e_* the elementary charge, and *Vol* the volume of a cell (sphere of 10 µm diameter). Calcium influx was also dependent on opening of channelrhodopsin given the fractional permeability *α* of the opsin to Ca^2+^ (see values in table below). [Ca] was increased after a spike by *Ca_influx_* = 1 µmol·L^−1^. The accumulated calcium then activated an afterhyperpolarization (AHP) conductance, *g_AHP_*, of 0.015 µS that reversed at *E_K_* = −80 mV, thereby producing the calcium-dependent AHP current *I_AHP_* = *g_AHP_* · [*Ca*] · (*E_K_* − *V*).
- An independent Ornstein-Uhlenbeck current noise *I_noise_* was added to each LIF neuron in order to capture intrinsic fluctuations in membrane potential. This noise was generated as a mean-reverting process with rate constant θ = 2 s^−1^ and noise intensity σ = 0.1 s^−1/2^. With these different current sources, the membrane potential *V_m_*(*t*) of the SGN obeyed:

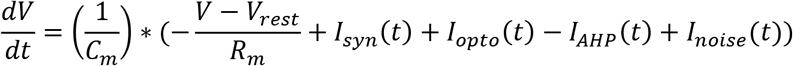

Intrinsic properties of LIF neurons were drawn from Gaussian distributions (σ = 10%) to account for cellular variability (Table 1).

**Table 1.** Intrinsic properties of High, Medium, and Low Spontaneous Rate fibers (HSR, MSR, LSR).

|  | HSR | MSR | LSR |
| --- | --- | --- | --- |
| $R_m$ ($\text{M}\Omega$) | $100 \pm 10$ | $90 \pm 9$ | $80 \pm 8$ |
| $C_m$ (pF) | 8 | 7.5 | 7 |

#### Parameter optimization with sound stimulation

SGN were grouped by spontaneous rate (low (LSR), medium (MSR), and high (HSR)) with 10 neurons of each SR per CF (i.e. 30 neurons/CF). Vesicle-release dynamics followed parameters from ref. (36) except for: 1) the immediately releasable pool capacity M = 10 vesicles, which was decreased to avoid multiple spike events and 2) the replenishment rate x = 500 s⁻¹ and re-uptake rate r = 20 s⁻¹, which were increased to avoid too strong adaptation and match SGN dynamics as seen in biological experiments (76, 77). Parameters affecting vesicle release rate (calcium time constant in seconds *τ_SR_* = [25*e*^−6^, 75*e*^−6^, 130*e*^−6^] and calcium conversion factor *z* = [10*e*^42^, 2.3*e*^42^, 2.3*e*^42^]) were optimized by mean-square-error optimization and further hand-tuned so that model rate-level functions matched experimental data (18).

Synaptic input to each LIF unit was generated by convolving synaptic “fusion events” with an exponential kernel of amplitude *I_syn_* = 0.5 nA and time constant *τ* = 1 ms. These values were hand-tuned to evoke consistent stimulus-evoked response without saturating the LIF unit, and a membrane time-to-peak consistent with *in vitro* whole-cell recordings in IHC afferents (49, 73).

Intracellular calcium was modelled with a single-exponential decay (*τ_Ca_* = 5 ms) and a fixed per-spike increment (*α_Ca_* = 1, in arbitrary concentration units). *τ_Ca_* was selected to reproduce the fast recovery of Ca^2+^-activated potassium currents; *α_Ca_* was then manually adjusted so that spike trains at 100 Hz elevated Ca^2+^ just enough to evoke realistic adaptation under continuous stimulation.

Finally, central connectivity was arranged in two successive chopper-unit stages with 30 neurons per channel. For each SGN type (LSR, MSR, HSR), 10 neurons converged onto 1 CN chopper neuron. The spatial distribution of the inputs was randomly drawn with a Gaussian width σ = 0.5 (in units of channels). For each CF and SR, CN units then projected to IC chopper neurons, again using a 0.5 Gaussian input profile (10 CN inputs per IC neuron). This σ = 0.5 spread was chosen so that each chopper neuron receives the bulk (between 7-8) of its inputs from its own CF channel, with a graded fall-off to neighboring channels.

#### Optical stimulation

To mimic biological experiments, we varied both the illumination level and the repetition rate of light pulses. For variable light intensity, we delivered a single 1 ms rectangular light pulse with irradiances logarithmically spaced between 0.1 and 1000 mW·mm^−2^ (30 values). For variable train rates, we presented 100 ms trains of 1 ms pulses at rates of 20, 30, 40, 50, 60, 80, 100, 150, 200, 250, 333, or 500 Hz with a fixed irradiance of 50 mW·mm^−2^. Each stimulation was repeated in 50 independent trials.

Spatially, the light was assumed to illuminate all 21 characteristic-frequency (CF) channels with a Gaussian profile (σ = 5 channels) centered on the central channel (#11). The profile was chosen so that 50% of the channels received at least 50% of the maximum irradiance. To mimic biological experiments with stimulation of either IHC or SGN, the opsin conductance was inserted into the membrane potential equation of either IHC or of SGN.

Throughout all optogenetic simulations, the acoustic channel was silenced and feedback loops acting on cochlear amplification, both acoustic reflex and medial olivocochlear loop, were disabled. As a result, the middle ear and dual-resonance nonlinear filter bank stages were not sending any input to the IHC stage, whose synaptic process was still operant.

#### Parameter optimization of optogenetic stimulation

For wild-type ChR2 we adopted all transition rates unchanged and hand-tuned only the maximal conductance *G_max_* so that a 100 ms, 10 mW·mm^−2^ pulse produced 450 pA peak and 26 mV depolarization as in our patch-clamp calibration experiments (Fig. S1). For the red-shifted variants both the rate constants and *G_max_* were adjusted to match published kinetics and amplitudes (38, 74). For SGN stimulations, G*_max_* was fixed at 0.011 µS in order to evoke ∼1 spikes for a 1 ms light pulse at 50 mW·mm^−2^, as seen in experiments from others (38, 74).

Finally, light-gated calcium influx was set to equal 5% of the ChR2 conductance (78), scaled by an inward-rectification factor and physical conversion constants (79). This fraction was varied in order to match adaptation in firing upon repetitive light stimulations.

#### Numerical simulations

All simulations reported in the present paper were run with the settings and parameters summarized below (Table. 2 and 3). Spikes of SGN, CN and IC stages were saved for further analyses as in experiments.

**Table 2.** General parameters of the auditory model.

|  |  |
| --- | --- |
| Sampling Frequency | 100 kHz |
| Segment Length | 10 ms |
| Spatial profile of optical stimulation | 5 channels |
| Central connectivity | 0.5 channel |
| Vesicle stock M | 10 |
| Replenishment rate | 500 s<sup>-1</sup> |
| Re-uptake rate | 20 s<sup>-1</sup> |
| $\tau_{SR}$ (LSR, MSR, HSR) | [25, 75, 130] μs |
| Calcium Conversion factor z (LSR, MSR, HSR) | [10, 2.3, 2.3] $\cdot 10^{42}$ |
| $I_{syn}$ | 0.5 nA |
| $\tau_{syn}$ | 1 ms |
| $[Ca]_{influx}$ | 1 μmol·L<sup>-1</sup> |
| $\tau_{Ca}$ | 5 ms |
| $g_{AHP}$ | 0.015 μS |

**Table 3.** Parameters of the optogenetics current.

|  |  |
| --- | --- |
| $\alpha_{ChR2}$ | 0.5 % |
| $\alpha_{fChrimson}$ | 0.025 % |
| $\alpha_{fChrimson\_noCa}$ | 0 % |
| $g_{ChR2}$ | 0.027 for IHC, 0.011 for SGN in $\text{mS}\cdot\text{cm}^{-2}$ |
| For f-Chrimson variants: |  |
| $e_{12,dark}$ | $0.022 \text{ ms}^{-1}$ |
| $e_{21,dark}$ | $0.016 \text{ ms}^{-1}$ |
| $G_{d,2}$ | $0.1 \text{ ms}^{-1}$ |
| $\epsilon_1$ | $2 \text{ ms}^{-1}$ |
| $\epsilon_2$ | $0.2 \text{ ms}^{-1}$ |
| $G_r$ | $2*0.0000434587*\exp(-0.0211539274*V)$ |
| $G_{d,1}$ | $2 * (0.075 + 0.043.* \tanh((V + 20)/-20))$ |

#### Simulation and decoding of a sound library

We used the <u>ESC-50</u> dataset that contains 2,000 labelled environmental audio recordings. The dataset consists in 5 s long recordings organized into 50 semantical classes. Each class contains 40 examples of sounds that can be further grouped into 5 major categories (animals, natural soundscapes and water sounds, Human non-speech sounds, interior/domestic sounds, exterior/urban noises).

For acoustic stimulation, we used the full model and adjusted the maximal sound intensity during the 5 s epoch to 90 dB. For optogenetic stimulation, each individual sound was first transformed into a spectrogram. The spectrogram was divided into 21 frequency bands by filtering the sound stimulus with a bandpass Butterworth filter of order 6, of central frequencies logarithmically distributed between 250 Hz and 8 kHz, and of frequency bandwidth at 3 dB equal to the spacing between 2 consecutive central frequencies. The power of each frequency band was lowpass filtered at 400 Hz and Hilbert transformed to extract the envelope. The envelope was passed through a logistic function (amplitude: 1; steepness: 15; midpoint: 0.4) and every value below 0.05 was set to 0 to avoid constant light stimulation. The resulting waveform was used as the stimulation light intensity and its amplitude was scaled such that a pure tone of intensity 90 dB would generate a light intensity of 50 mW·mm^−2^.

After building the different stimuli, spiking activity in the SGN was computed by running the different models (acoustic, optogenetic stimulation of IHC or SGN with different opsin variants) with a sampling rate of 44.1 kHz. Neurons from the same type (either HSR, MSR, or LSR) and from the same frequency channels were binned together, resulting in 63 ‘neurons’ (3 types × 21 channels). Temporal binning was also used in the temporal domain with a window of 512 data points (11.6 ms). Thus, the generated data (630 neurons × 220500 timesteps) was reduced to a matrix of size (63 × 430). Each matrix was further segmented into 5 consecutive sections of size (63 × 86).

To perform audio classification on the ESC-50 dataset, we implemented a convolutional neural network (CNN, Fig. S13) operating on time-frequency representations of the spiking data (neurogram) similar to previously developed CNN working on audio spectrogram (39). The input to the network was a segment of the neurogram with dimensions (63 × 86). The model consisted of two convolutional blocks followed by three fully connected layers. The first convolutional layer used 80 filters with a kernel size of (60 × 6), followed by batch normalization, a ReLU activation function, and max-pooling with a kernel size of (4 × 3) and stride (1 × 3). The second convolutional layer also contained 80 filters, using a (1 × 3) kernel, followed by batch normalization, ReLU activation, and a second max-pooling operation with a kernel size of (1 × 3) and stride (1 × 3). The resulting feature maps were flattened and passed through two fully connected layers, each containing 5000 neurons with ReLU activation. Dropout with a probability of 0.5 was applied after each fully connected layer (FC) to reduce overfitting. The final layer was a fully connected classification layer with (*N_classes_*) output units, where (*N_classes_*) corresponds to the number of target sound classes. The network output logits were used for multi-class classification.

To improve classification robustness, each 5 s recording from the ESC-50 dataset was split into five consecutive 1 s segments. The decoder classified each segment independently, generating a probability distribution over the 50 sound categories. The final prediction was obtained using probability-weighted voting across the five segments, reducing the influence of isolated classification errors and better capturing information distributed throughout the recording (39).

**Figure S13.**
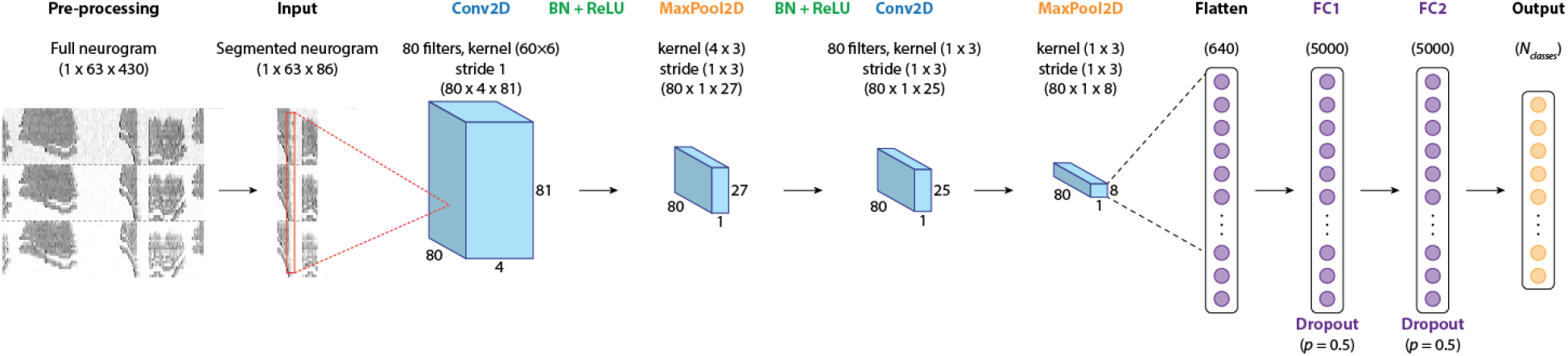
Architecture of the convolutional neural network. See Material and Methods for detailed explanation.

The large computational cost of simulating all 2,000 sounds across multiple stimulation modes and opsin variants was addressed through code optimization, parallelization, and deployment on an HPC cluster using SLURM, reducing simulation time from over 25 hours to approximately 1.5 hours per configuration. Decoder training was also accelerated on NVIDIA A40 GPUs, allowing the complete evaluation pipeline (simulation, training, and validation) to be completed in less than one day instead of nearly one week.

## Author Contributions

JB designed, formally supervised the project and provided funding. WL, ASP, and JL performed the experiments. WL, VB and JB analyzed the data. VB performed the simulations. ASP and TP performed the histology and imaging experiments. ASP, VB, TP and JB wrote the manuscript.

## Data, code, and materials availability

The datasets generated and analyzed during the present study will be available on the Institut Pasteur storage center. Data acquisition (Labview), analysis (Matlab), and simulation (Matlab) softwares used in this paper are described in the Methods and will be available upon reasonable request.

